# Identifying reliable fitness proxies for growing animals responding to anthropogenic changes

**DOI:** 10.1101/2020.08.06.239616

**Authors:** Andrew D. Higginson

## Abstract

Anthropogenic influences on habitats often affect predation on species by introducing novel predators, supporting additional predators, or reducing animals’ ability to detect or avoid predators. Other changes may reduce the ability of animals to feed, or alter their energy use. An increase in predation risk is assumed to reduce prey populations by increasing mortality, reducing foraging and growth. Often animals don’t appear to have been adversely affected, or may even increase growth rate. However, theoretical predictions that may have been overlooked suggest that optimal foraging rate, mortality rate and growth rate may change in counter-intuitive ways, depending on exactly how predation risk or costs have been increased. Increasing predator density may increase mortality rate when foraging, reduce the safety of refuges, or alter the relationship between vigilance and attack likelihood. Increasing temperature may increase metabolic costs in ectotherms and reduce thermogenesis costs in endotherms, which affects the costs of active foraging and inactivity differently. Here, I review the theory on how predation risk and metabolic costs should affect foraging behaviour, mortality and growth in order to explain the great variation in behavioural responses. I show that in some situations animals may not respond behaviourally even though a change severely affects survival, and the mortality may be a poor metric of the impact of a change on population viability. In other situations a fitness proxy may change dramatically whilst fitness is unaffected due to compensatory changes in behaviour or life history. Other measures may change in a positive way whilst fitness declines. I describe how to identify the situations in the field and thereby make reliable measure of fitness in particular study systems. Overall, this work shows how behavioural theory can help understand the impacts of environmental change and highlights promising directions to better understand and mitigate their effects on ecosystems.

## 0. INTRODUCTION

Anthropogenic influences on habitats often affect predation on species by introducing novel predators, supporting additional predators, or reducing animals’ ability to detect or avoid predators (Sih et al., 2011). An increase in predation risk is often assumed to reduce prey populations by increasing mortality, but also reducing foraging and growth due to behavioural responses that reduce predation rate (Creel et al., 2019; Sheriff et al., 2020; Sinclair et al., 2008). The non-lethal effect of predation risk on ecosystem stability may be more important than the mortality (Lima & Dill, 1990) because prey responses may alter their survival or reproduction with subsequent effects on other species (Bolnick et al., 2011; Lima, 1998). It is therefore crucial to understand how anthropogenic changes alter behaviour and reproductive success.

It is commonly assumed that increasing the predation rate, such as by increasing the number of predators or risk in experiments, will reduce the foraging rate and so prey growth rate (Suzuki et al., 2010). Predator presence cause an increase in vigilance (and so reduction in foraging), although there was variation between prey species (Creel et al., 2019) and within prey species (Ramamonjisoa et al., 2019). Sometimes the opposite effect is observed (Dmitriew, 2011; Preisser & Orrock, 2012; Verdolin, 2006) and there is no correlation between the strength of antipredator responses and direct predation (Creel et al., 2017). This may indicate differences in the trade-off between gaining nutrients and vulnerability to predation (Creel, 2011).

An important environment pollutant that affects behaviour is anthropogenic noise, such as from shipping. A meta-analysis of the effect of aquatic noise found a wide range of behavioural responses, from very negative reductions in foraging to positive effects (Cox et al., 2018). In terrestrial species noise affects zebra finch foraging rate (Evans et al., 2018) and may increase risk by reducing the effectiveness of alarm calls in cooperatively foraging groups (Shannon et al., 2020). Some studies have managed to measure not just foraging or growth but mortality, and there are strong effects of boat noise on the mortality of reef fish (Simpson et al., 2016). Other anthropogenic changes include the turbidity of water caused by sedimentation or eutrophication, which increases risk and alters foraging behaviour (Figueiredo et al., 2020), and disturbance by people can make antipredator responses more rapid (Van de Voorde et al., 2015). Temperature change caused by climate change could have profound effects on predator-prey dynamics (DeLong & Lyon, 2020) via effects on activity rate and/or metabolic energy use (Johansen et al., 2014). However, a number of experiments have failed to find an effect of noise on mortality (Wysocki et al., 2007) and growth (Bruintjes & Radford, 2014; Wysocki et al., 2007), with one finding that noise pollution delays maturation but does not affect reproductive output (Gurule‐Small & Tinghitella, 2019).

Environmental factors affect the strength of antipredator behaviour (Moll et al., 2017). The landscape structure is highly important because the provision of refuges affects the effectiveness of responses to predation risk (Heithaus et al., 2009). Predators prefer to forage where there is less cover for prey (Haram et al., 2018) whilst prey select locations with more refuges (Bishop & Byers, 2015). The availability of food in locations that are safe from predators have strong impacts on growth and reproduction in mice (Arthur et al., 2004) and fish (Crowder & Cooper, 1982), and refuge from predation also affect the timing of life history decisions such as maturation (Grof-Tisza et al., 2015). Habitat changes such as loss of complex vegetation and invasive plants may alter the availability of refuges and so the vulnerability of prey to predation (Norbury & Overmeire, 2019).

Researchers are typically interested in the impact of changes on animals fitness because this affects the viability of populations. However, reproductive success is challenging to measure and typically studies measure some proxy of fitness, such as growth rate, mortality, or survival. Ultimately, these studies seek to understand how changes will affect the viability of populations, so ideally measure population size change, which can be measured directly or estimated from measures of individual lifetime reproductive success. The latter is the product of the probability of surviving to maturation and the reproductive success if maturation is reached. It may be challenging to measure both of these, either because only adults can be observed or because offspring production cannot be measured. Sometimes is not feasible to measure either, so some other proxy of fitness such as mortality, growth, or foraging rate is measured.

Animals trade off predation against growth, but also survival to maturation (decreasing with adult body size) against the reproductive success after maturation (increasing with adult body size),(English et al., 2016; Houston & McNamara, 2014). Hence, measuring a trait such as growth rate early in development to use as proxy of fitness may not capture the full consequences for a population. Models of foraging behaviour show that animals may not respond in ways that we might expect. For instance, the adaptive response to a reduced ability to detect predators, which will increase predation risk, may result in increased foraging and faster growth under certain conditions (Houston & McNamara, 1989; McNamara & Houston, 1987). No effect of noise on growth may not mean that the noise will not negatively impact an ecosystem if it affects survival. On the other hand, different behaviours and life history traits tend to adjust in concert and to a greater or lesser extent cancel out one another’s effects on adult fitness. Hence, a seemingly dramatic decrease in early growth may be taken to imply a dramatic decrease in adult fitness, but survival may be increased so the effect on adult fitness could be negligible.

Understanding these responses are critical to predicting effects of changes on ecosystems, as behavioural plasticity can be a quick route to evolutionary rescue (Carlson et al., 2014) that enable species to persist in a changed world (Sih et al., 2011). It is crucial to use theoretical models to interpret the results of experiments and direct future experiments. However, the theoretical literature is often somewhat opaque (Fawcett & Higginson, 2012) so some models may not be known – or their implications not clear – to behavioural ecologists and conservation biologists. Here, I review some models that have captured the trade-off between growth and predation in the hope of explaining their usefulness for understanding the responses of animals to alterations to their environments in predator density, food availability, refuge availability, or ambient temperature.

## 1. MINIMISATION OF THE MORTALITY RATE TO GROWTH RATE

For growing animals under the risk of predation it can be assumed that they act to minimise the ratio of the mortality rate to the growth rate (Gilliam, 1982; Werner & Gilliam, 1984). For instance, if an animal has to reach a certain size to mature then minimising this ratio will maximise the probability of surviving to maturity. The situation becomes complex if some other state changes as they grow, such as investment in antipredator defences (Higginson & Ruxton, 2010), if the food supply is variable and they can store nutrients (Higginson et al., 2012), or if there are strong time constraints (Houston et al., 1993). I will not address these complexities here because there is sufficient complexity that is not widely acknowledged just in how activity affects the rates of growth and mortality.

Animals may not choose the option that has the highest growth rate nor the one with the lowest mortality rate, depending on the form of the trade-off. Consider there are a range of options controlled by the decision *x* that affects both mortality and gain, such as foraging rate (e.g. searches per unit time). Assume that every search has fixed probability *r* of resulting in a food item so the gain rate is *rx*. Consider that the predation rate increases as foraging intensity increases *p*_*1x*_, and there is another source of mortality of rate *p*_2_. Then the mortality to gain ratio is

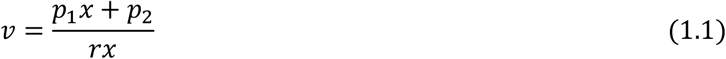

To find where the minimum is we try to differentiate with respect to foraging rate *x* to find where the slope is zero.

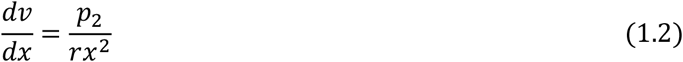

Since all parameters are positive this is always positive: *v* always increases as *x* increases, so foraging should be minimised. Researchers have usually assumed that predation rate increases in an accelerating way as foraging increases, perhaps because no vigilance is much worse than a little vigilance, but vigilance has diminishing returns (see Appendix A). Alternatively, animals may first forage in safer places and only in dangerous places if they forage at a very high rate. Whatever the justification, this assumption means that there can be a solution to the equation, provided there is some other complexity such as a source of mortality.

Response to predation risk depends on how activity affects mortality rate (McNamara & Houston 1993). If there is no other source of mortality the value ratio is

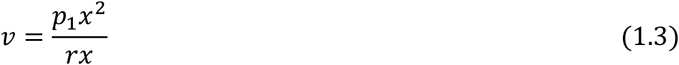

then the slope is

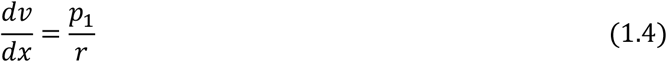

which since all parameters are positive is also positive, so foraging should again be minimised (Figure 1a). If there is another source of mortality with rate *p*_*2*_ then

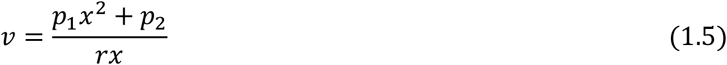

then as the predator density *p*_*1*_increases the optimal foraging rate decreases (Figure 1b).

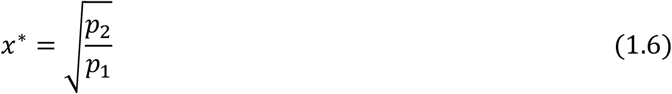

**Figure 1:**
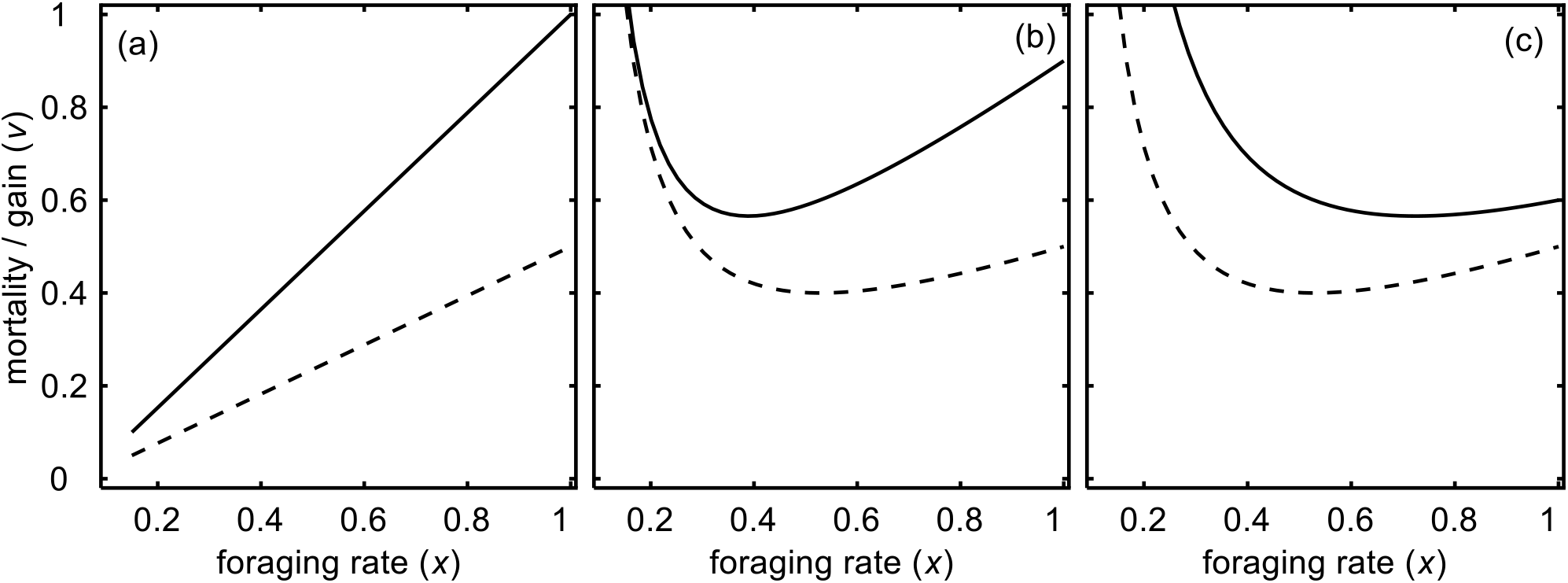
Effects of foraging rate (*x*) on the ratio of the mortality to gain rates (*v*) for a doubling of the risk of predation *p*_*1*_. (a) *r* = 1.0, *p*_*2*_ = 0, *p*_*1*_ = 0.5, 1.0; (b) *r* = 1.0, *p*_*2*_ = 0.1, *p*_*1*_ = 0.4, 0.8; (a) *r* = 1.0, *p*_*2*_ = 0.1, *p*_*1*_ = 0.1, 0.2. Dashed lines are for smaller *p*_*1*_ values; solid lines for larger *p*_*1*_ values. The optimal foraging rate (*x\**) is where the value ratio (*v*) is minimised.

If however the predator density does not interact with foraging rate, but foraging rate affects the other source of mortality, such as by exposing the animal to pathogens, the value ratio is

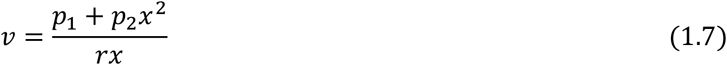

then if the predator density increases the optimal foraging rate *increases* (Figure 1c).

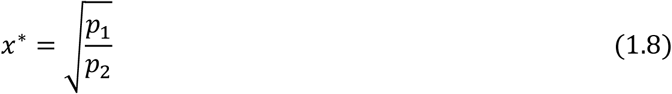

In summary, the response of prey animals to increased predation risk may be positive, negative, or neither depending on the relative size of the effects of foraging and not foraging on mortality rate. It is therefore important to have some idea of the interaction between foraging and mortality rates. Below, I explore this further with more realistic details.

## 2. EFFECT OF PREDATOR DENSITY ON THE GROWTH-MORTALITY TRADE-OFF

Now I assume that there is curvature in the effect of foraging on predation risk by raising to the power 1/σ (I use the reciprocal because it makes the presentation simpler below). The ratio of mortality to growth (net gain) rates is

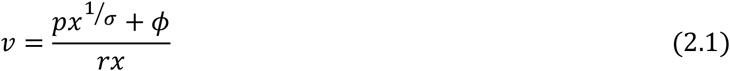

An increase in exposure to predators due to environmental change or in an experiment may affect the mortality function in one of three ways: increase in overall risk *p*, increase in baseline risk *ϕ*, and/or increased risk at intermediate foraging rates such as if vigilance becomes less effective, which may not affect risk at constant vigilance or constant foraging but the trade-off between vigilance and foraging (σ, see Figure 2a). Each of these alters the mortality-growth trade-off in different ways (Figure 2b). The optimal foraging rate does not depend on *r* but changes in different ways when the predation risk parameters increase (Figure 2c).

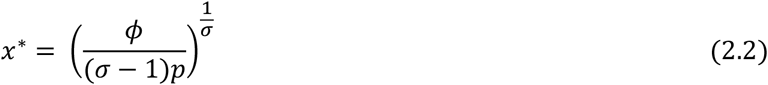

**Figure 2:**
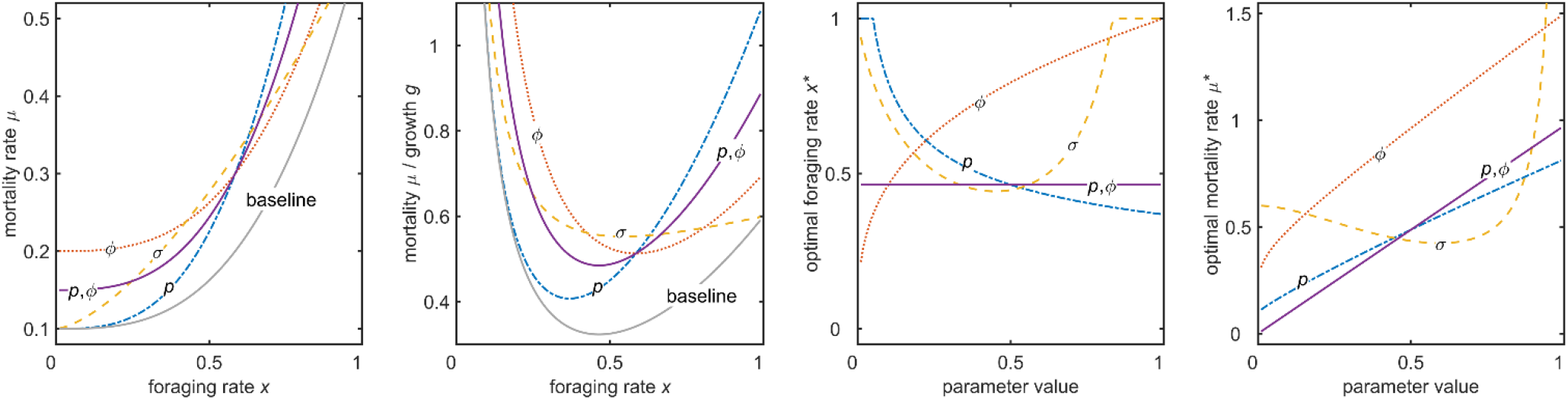
Plausible effects on the relationship between mortality rate *μ* and foraging rate *x* and the optimal foraging rate when exposure to predators increases. The predator density may affect the overall risk *p*, the relative risk when inactive ϕ, or the trade-off between foraging and avoiding predation σ, or a combination (e.g. both *p* and ϕ). Baseline parameter values are *p*=0.7, ϕ=0.2, σ=2. (a, b) Each line is labelled by the parameter that was increased (*p*=1.0, ϕ=0.4, σ=1.5). The optimal foraging rate is the value of *x* where the curves are at the minima (c) Change in optimal foraging when each parameter increases from 0 to 1, (d) Change in mortality rate for optimal *x* when each parameter increases from 0 to 1.

Firstly, note that if ϕ is a fixed proportion of *p* (e.g. *k*/*p*) then there is no effect of *p* because *p* would be in both the numerator and denominator so would cancel out (solid lines in Figure 2). This counter-intuitive result occurs because it is the *relative* risks that matter, not the absolute values. Thus, despite the fact that increased predator density increases mortality (Figure 2d) the animal should not respond to it. Hence a lack of behavioural response does not indicate that an environmental change will not influence prey populations.

If the baseline risk increases (ϕ increases) then foraging rate increases, because if the animal can do nothing about risk then they should try to reach maturity as quickly as possible. In instead the increase in risk is linked to foraging (*p* increases) then the animal should reduce foraging. On the other hand, the effect of the curvature (σ) is u-shaped, so foraging rate may increase or decrease depending on the extent to which foraging already affects the ability to avoid predators (i.e. the shape of the curve in Figure 2a).

As a result of these differences in the foraging response the dependence of mortality rate on mortality risk has different shapes in the three cases (Figure 2d). It is again u-shaped as σ varies. When σ is large the mortality rate is flat over most of the range of *x* and the foraging rate is optimal just left of the steep increase in mortality. This point is at smaller *x* as *σ* increases. Above a certain value of σ the trade-off becomes so strong (almost linear, Figure 2a) that intermediate *x* are no longer advantageous and the animals should forage at full rate.

## 3. EFFECT OF BENEFITS AND COSTS WHEN NOT FORAGING

The above analysis shows that the important aspect of increasing predation risk on behaviour is not overall risk, but how much prey can affect risk. Similarly the effect of the animal on its metabolic costs and the foraging gain are important. It is therefore crucial to consider what happens when the prey animal is not foraging (usually assumed to be when obviously searching), even though prey that are not active may be able to able to eat, may still experience a risk of mortality and will still have metabolic energy needs. In this section, I show that such costs can strongly affect predicted behaviour.

Assume that an animal spends *x* proportion of its time actively foraging (0≤*x*≤1) and the rest of the time immobile in a refuge, scanning for predators, waiting in ambush for prey, or another activity. This activity is less costly in terms of energy use and has lower mortality risk, but results in smaller gain. For brevity I refer to this activity as “resting”.

Predation rate is again the probability per unit time of being killed is proportional to the predator density, increasing in an accelerating way with foraging rate and influenced by the relative risk when resting. For clarity in this section I assume that the exponent (σ) is 2.

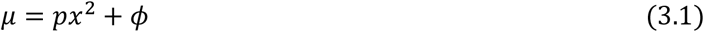

Energy gain rate is assumed to increase with foraging activity, food availability and the availability of food when resting

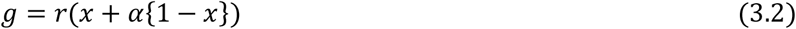

where *r* is food availability and α is the relative gain when resting (0≤α≤1).

Similarly, energy use rate is the sum of metabolic scaling multiplied by the cost of activity and the cost of resting (e.g. basal metabolic rate), which is a proportion of the cost of being active.

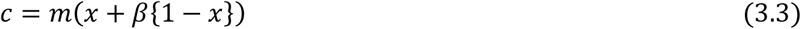

where *m* is metabolic costs and β is the relative cost when inactive (0≤β≤1).

Fitness again the ratio of the mortality to foraging rates

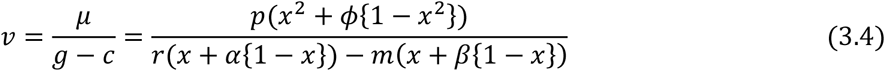

To find the optimal foraging rate we differentiate with respect to *x* and find where the slope is zero: this is where *v* is minimised. We can assume *r*=1 without loss of generality of the results because *m* can be imagined to be scaled to the maximum gross energy gain. So, an increase in food means a decrease in *m*, and an increase in metabolic costs means an increase in *m*. The optimal foraging rate is

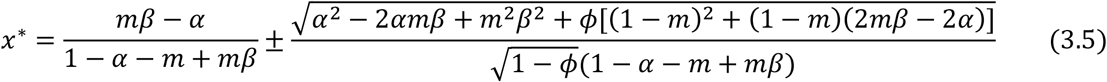

We can use *x\** to get the optimal mortality rate and the optimal growth rate by substituting (3.5) into the equations for *μ*(*x*) and for *g*(*x*) − *c*(*x*).

The effect of all parameters on foraging *x\**, mortality *μ*(*x\**) and growth *g*(*x\**) – *c*(*x\**) are shown in Figure C1. Optimal foraging rate does not depend on food availability *r* which is merely a multiplier so mortality is unchanged and growth increases (Figure C1g,m). Optimal foraging rate decreases as α increases, because it is less worthwhile foraging, so mortality declines but growth increases (Figure C1h, n). *x\** increases as *m* increases to try to maintain high growth which is only effective until maximum foraging rate is reached (Figure C1i). Increasing β causes *x\** to increase because the relative cost of activity is smaller, and so growth and mortality increase. As *p* increases *x\** decreases which means no increase in mortality but a steep decrease in growth rate (Figure C1k,q). ϕ has the opposite effect on foraging so growth increases but mortality increases very steeply (Figure C1l,r).

Since observations and experiments often focus on growth and mortality rates as a measure of the impact of environmental changes, I consider how they depend on the parameters (mathematical working in Appendix D). However, experiments will be designed to detect a difference between a treatment and control group. In Figure 3 I show where the effect of each parameter is small enough to be considered as no effect. Figure 3a-d (more parameter space shown in Figure C2) shows the effect of the parameters (different columns) on the foraging rate, where cyan indicates small change, blue a positive effect and green a negative effect (black indicates foraging is at maximum rate and white indicates at minimum rate so there is no change). Optimal foraging rate mostly changes as expected: negative if *p* or α increases and positive if *m* or β increases. The exception is the effect of *m* where β<α (bottom of Figure 3b) because an increase in *m* has a smaller effect on energy use when resting than the benefit of activity so it is optimal to forage less.

**Figure 3:**
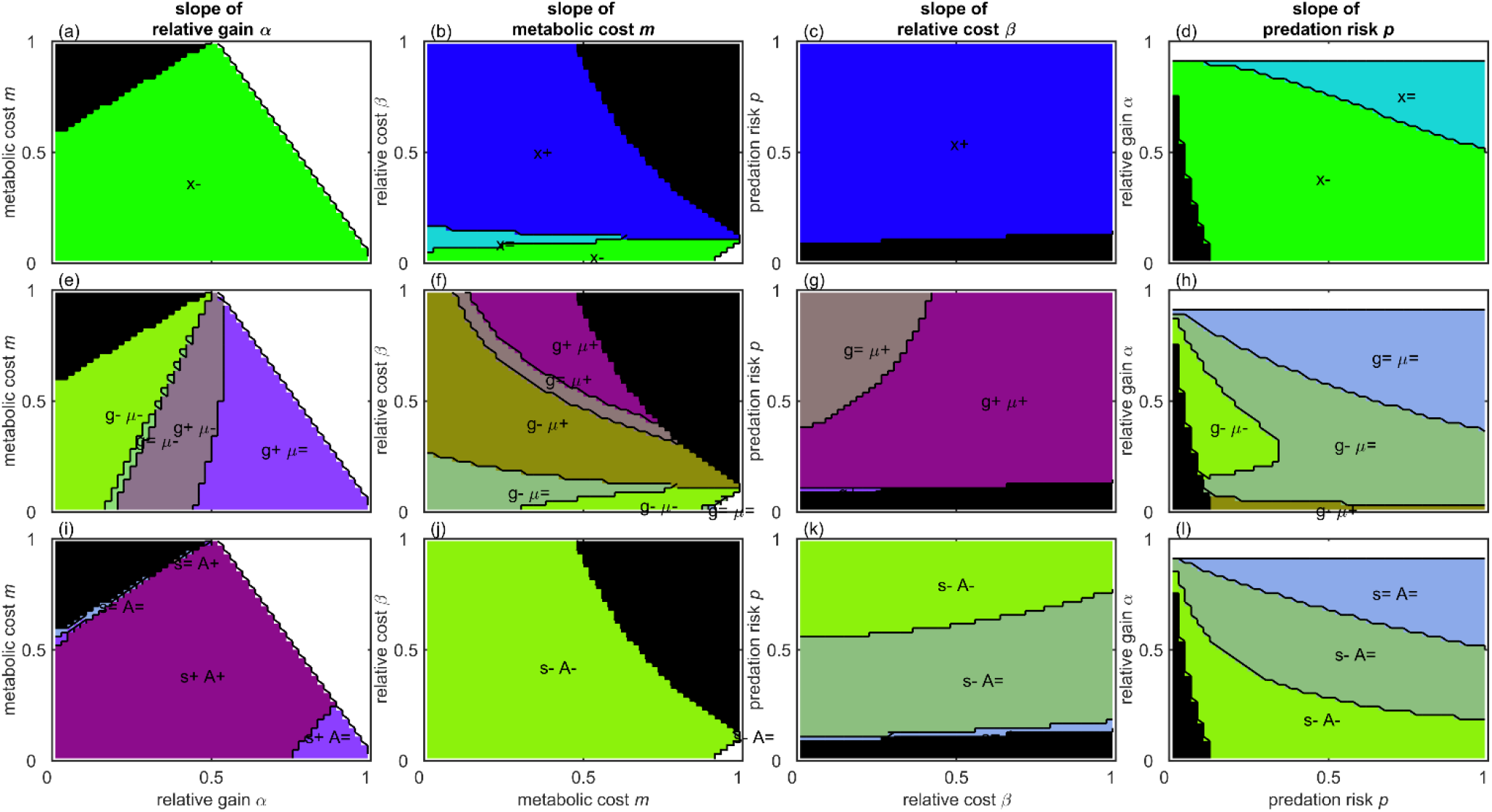
Effect of each parameter (columns) for the values of the parameter (x-axes) and some of other parameters (y-axes) on (top-row, a-d) optimal foraging rate, (middle row, e-h) growth and mortality rates, (bottom row, i-l) survival to- and size at maturation. Black areas are where *x*≥*1 and white areas where *x*≤*0 so does not change.

Figure 3e-h (more parameter space shown in Figure C3) shows the change in growth and mortality rates. The change in the optimal growth and mortality rates are usually as expected: an increase in resources positively affects growth and negatively affects mortality, whereas an increase in costs positively affects mortality and negatively affects growth. However, the strength of these effects may be very small, and there are conditions where the opposite effects are predicted. Often the critical thresholds for the opposite effects are where *x\** is out of range (*x\**>0 or *x\**<1). There would be no change in foraging (stay either minimal or maximal) so the effect would be as expected without a response: α positively affects growth, *m* and β negatively affect growth, *p* and ϕ positively affect mortality. However, there are many interesting exceptions.

### Effect of free food

Within the range 0≤*x*≤*1 increasing α always reduces mortality although the effect is very small if α is large because the change in *x* is small (purple area). Increasing α may decrease the growth rate if free food was not available (Figure C3a) because animals reduce foraging rate. A change in the relative gain when resting (α) can have positive (purple) or negative (green) effects on growth rate depending on its current magnitude. If α is large then activity is low, so an increase in α simply increases gain rate. On the other hand, if α is low then activity is high, so an increase in α makes it beneficial to reduce activity strongly to reduce predation but without reducing gain too much, so growth rate decreases. This range of activity changes are the reason for the great difference in the magnitude of the change in mortality rates, from strongly negative to negligible across the range of α. The critical values of α depend on the other parameters, but this pattern is broadly unchanged (Figure C3a, f, k, p).

### Effect of ratio of metabolic costs to food availability

A change in the metabolic costs *m* usually results in reduction in growth and increase in mortality, but the magnitude of these effects strongly depends on other parameters (not so much on itself, Figure C3): Figure C3f shows the effect of the relative cost when inactive (β). When *m* increases there is an *decrease* in mortality if β < α and an increase in growth under some conditions (condition D4), also when β is large. When β is large (resting almost costly as costly activity) activity increases in response to an increase in *m*, so both growth and mortality rates increase. When β is small the optimal response is to reduce activity or only increase it slightly, so growth rate decreases and mortality increases only negligibly.

### Effect of relative cost of resting

An increase in β might be expected to have negative effects on growth and a positive effect on mortality, but usually has a positive effect on growth because the animal should increase foraging rate. However, the magnitude varies greatly again. β and *p* interact to determine the effect of β (Figure 3g). When β is small and *p* is large the increase in foraging is slight so the effect on growth and mortality would be negligible (brown region). However, this does not mean that the effect on fitness is negligible, as when *p* is large the activity rate is low even if resting is equally costly as activity (β=1) so the effect of β on fitness is large.

### Effect of predation risk

An increase in *p* increases growth if *m*(1 − *β*) < 1 − *α*, but this latter is when *x*\*>1 so growth would be unaffected by *p*. An increase in *p* decreases mortality if α > *mβ* because the animal greatly reduces foraging. An increase in predation risk *p* leads to significantly increased mortality and decreased growth only in small region of parameter space (Figure C3e,j,o,t). When metabolic costs are small compared to food availability (*m*<0.4) and α<0.5 the strong reduction in foraging rate means that mortality does not increase strongly, but instead growth rate strongly decreases. This is the case in most of parameter space (Figure C3). When *m* is greater the animal cannot afford to reduce foraging very much, so mortality does increase too.

## 4. PREDICTING EFFECTS OF ENVIRONMENTAL CHANGE ON ADULT SIZE AND FITNESS

We can use the method of English et al (English et al., 2016) to find the optimal size of maturation and so the survival to maturation, payoff when mature, and overall fitness of an animal in response to any parameter (see Appendix B for the general approach). Here, I assume that the fitness after maturation *w* is simply the square root of body size *s* minus some minimum size for reproduction (ψ),

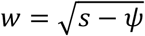

but any function with diminishing returns would give the same results. The results (Appendix D) are more intuitive in that benefits (α) always increase survival and size and costs (*m*, β, *p*, ϕ) always decrease them, but again the magnitude of these effects varies (Figure 3, C4). An increase in α generally increases maturation size and survival, except where α is large where activity is already so low that the effect on survival is negligible (Figure 3i, C4a,f,k,p). An increase in *m* causes a significant reduction in both size and survival over almost all of parameter space (Figure 3j, C4b,g,l,q). By contrast, an increase in β only reduces size not survival as the behavioural response is smaller unless *p* is large (Figure 3k) or *m* is large and β is small (Figure C4h). An increase in *p* reduces both size and survival, but in much of parameter space the decrease in survival is slight (Figure C4e,j,o,t), especially where activity is already low (Figure 3).

## 5. PREDICTING THE RESULT OF CHANGES IN PREDATION RISK AND TEMPERATURE

Anthropogenic changes to environments often include alteration in predator risk, either by increase in the predator density or increase in the vulnerability to existing predators. For example, underwater noise pollution may reduce the ability of prey to detect and avoid their predators and denuded vegetation may reduce the availability of cover so making prey more conspicuous. It is commonly assumed that increased predation risk will reduce foraging, growth and survival. However, the above models show how important it is to consider exactly *how* the predation risk increases. Denote the overall risk by *q*. It may be that the risk increases proportionally when foraging and resting and the *relative* risk when resting is constant (ϕ is a constant proportion of *p*),(Figure C5a). On the other, hand if there is a safe refuge increasing predator density would not strongly increase when resting (*ϕ* is constant),(Figure C5b). A possible intermediate is that the increasing predator density first impacts when foraging but then when predator density gets very high it becomes difficult to avoid them and so risk when resting increases more steeply (Figure C5c).

As may be anticipated from the previous sections, the different direction of change of *ϕ* as predation risk (respectively: increase, constant, decrease) means that the direction of change of foraging rate *x\** is different between these cases. If the proportion is constant then the foraging rate is unchanged (Figure 4a) and so the growth rate unchanged too (Figure 4c). The mortality rate increases linearly (Figure 4b) and the proportion of individuals maturing declines slightly (Figure 4e). If instead the risk when resting does not change the foraging and growth rate decrease (Figure 4a, c), mortality increases only slightly (Figure 4b, e). In the third case ϕ increases as predator density increases so the foraging and growth rate *increase*, the mortality rate increases steeply (Figure 4b, e).

**Figure 4:**
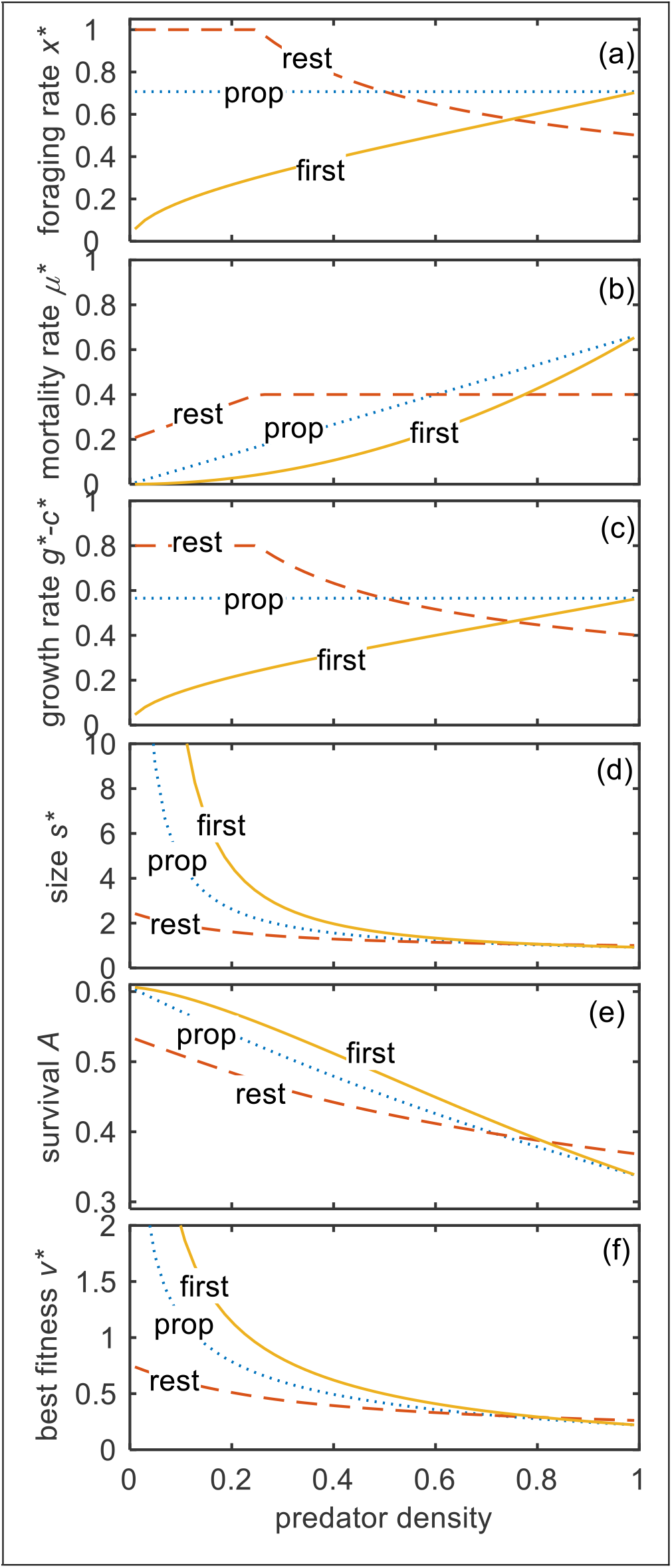
Predicted effect of increasing predator density (*q*) in experiments when the effect is one of the 3 in Figure C5 on (a) foraging rate *x\**, (b) mortality rate **μ*\**, (c) growth rates *g*–c\** (d) maturation size *s\** (e) survival to maturation *A* (f) fitness *w\**. Lines indicate three hypothetical effects of increasing predation risk: risk increases proportionally when foraging and resting (dotted lines); risk only increases when foraging and is constant when resting (dashed lines); risk increases first when foraging and then when resting (solid lines). Other parameter values *r*=1.0, α=0.1, *m*=0.1, *β*=0.5, *p*=0.4, *ϕ*=0.2.

Changes to climates may increase the ambient temperature in which animals live, which affects their metabolic costs. Poikilotherms may increase their metabolic energy use in warmer temperatures. Endotherms may be able to reduce their energy costs if the temperature increases, depending on how much heat is generated by foraging. Again, it could be the case that metabolic costs increase proportionally for both foraging and resting, so β is constant (Figure C5d). On the other hand, if the animals is an poikilotherm the cost of resting may increase more when resting if the temperature increases whereas active costs are constant (Figure C5e), meaning that β increases as temperature increases. A third possibility is that costs when resting decreases as the temperature increases if the animal is an endotherm in cold conditions and has to use thermogenesis when resting (Figure C5f), which may mean that the energy use whilst foraging is unchanged but the energy use when resting decreases. In each of these three possibilities β changes with different sign (constant, increase, decrease), as temperature increases, so the rates change in a different way (Figure 5). The foraging rate increases for constant total but decreases for proportional and constant difference and the mortality rate matches this. However, because of the change in costs the growth rate increases as temperature increases in the constant difference situation. In the constant total situation size, survival and fitness all increase (slightly) as temperature increases. In the other cases they decrease because costs are increasing.

**Figure 5:**
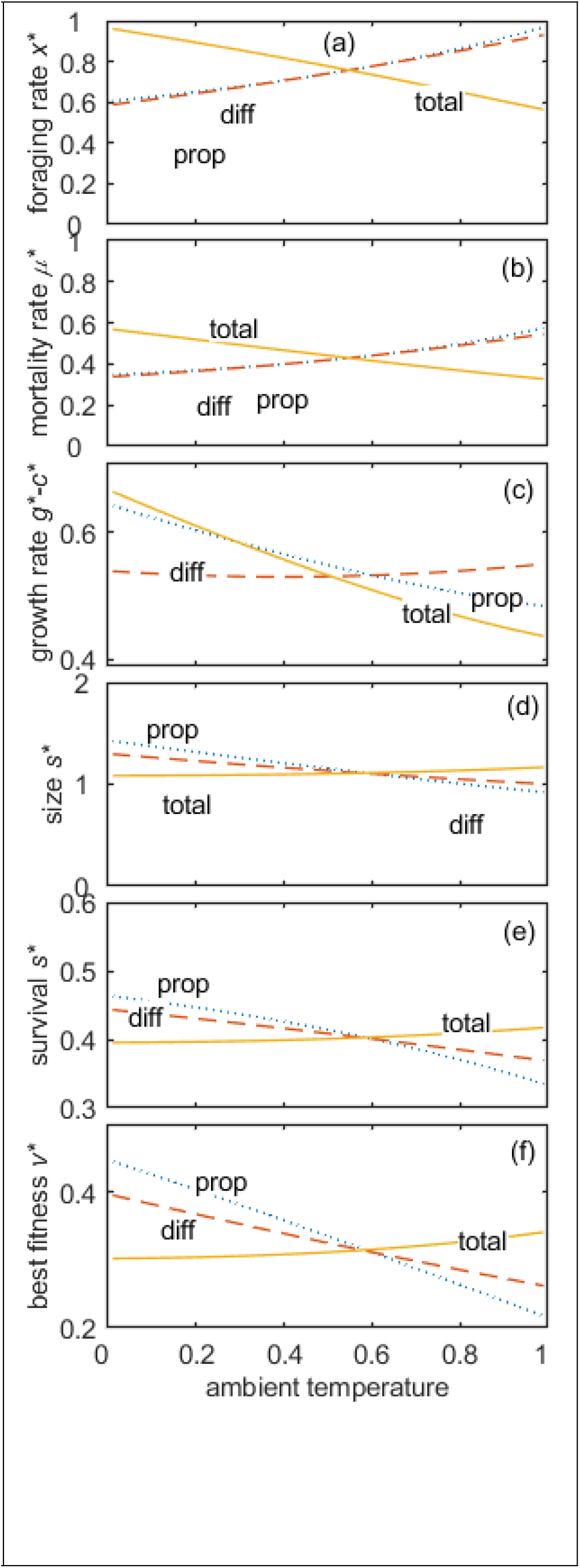
Predicted effect of increasing ambient temperature (*h*) in experiments when the effect is one of the three in Figure C5 on (a) foraging rate *x\**, (b) mortality rate **μ*\**, (c) growth rates *g*–c\** (d) maturation size *s\** (e) survival to maturation *A* (f) fitness *w\**. Lines indicate three hypothetical effects of increasing ambient temperature: metabolic costs increases proportionally when foraging and resting (dotted lines); costs only increases when resting and the effect of activity is constant (dashed lines); total cost of activity is constant and the cost of resting decreases (solid lines). Other parameter values *r*=1.0, α=0.1, *m*=0.1, *β*=0.5, *p*=0.4, *ϕ*=0.2

Changes in metabolic rate tend to be interpreted as maladaptive “stress”, but metabolic rate may change adaptively if it helps detect predators or evade attacks (Biro & Stamps, 2008; Careau et al., 2008; Houston, 2010). In Appendix E I show predictions for the optimal combination of metabolic rate and foraging rate when increasing metabolic rate decreases the predation rate. The analysis shows that the metabolic costs should not change if the baseline energy cost or the mortality risks change, so observed changes in metabolic energy use are likely to be non-adaptive.

## 6. DISCUSSION

Observations and experiments attempt to assess the impact of environmental changes on animals, such as disturbance, chronic noise, water turbidity, loss of vegetation, temperature, or reduced ability to capture food (Fletcher et al., 2012; Gurule‐Small & Tinghitella, 2019; Sih et al., 2011). In deciding how to respond animals will follow rules for trading off the benefits against the costs (Balaban-Feld et al., 2019; Dmitriew, 2011; Zimmer et al., 2011). Their choices may not be intuitive, and so the assumptions about the relationships between easily observed responses and the viability of populations may be incorrect. In reviewing theory of how animals trade-off growth against predation I hope to identify what proxies of fitness should be used in which situations in order to assess the impact of anthropogenic changes on populations.

I have shown that in some situations behavioural measures that are used to assess the impact of anthropogenic change on population viability may not be good measures. There are at least three possibilities: (1) The measure should not be expected to respond to the change, such as the foraging rate when predation risk increases both when active and inactive (Figure 4a,f); (2) The change is in the opposite way to fitness, such as when growth rate increases whilst fitness declines if the predation risk increases first when active before when inactive (Figure 4c,f); (3) The measure changes dramatically but fitness is barely affected, such as when the active costs are constant but the inactive costs decrease and the growth rate almost halves but the fitness is unchanged (Figure 5c,f).

In order to identify reliable proxies I calculated the strength of the association between each behavioural or life history measure and the fitness *w\** across the range of the environment variables in the six situations presented in Figures 4 and 5 (Figure 6). I added some stochasticity to the proxy and fitness measures and ran simple linear regressions to get *r*, the correlation coefficient between the proxy and fitness. The *r* values are around zero for a proportional increase in predation risk for foraging and growth rates (black bars), because they do not adaptively change (Figure 6a). The mortality rate is not a good proxy when the predation risk when resting is constant (grey bars). When the risk increases first when foraging the growth is negatively correlated with fitness (white bars). When temperature varies (Figure 6b) there is no good proxy for fitness in the constant total cost situation (white bars) because fitness varies very little, even though foraging, mortality and growth rates change two-fold over the studied range. Proxies are roughly equally good for proportional increase in costs (black bars) but there is a negative association between foraging rate and fitness. When the difference in cost is constant (grey bars) all proxies are reliable except the growth rate.

**Figure 6:**
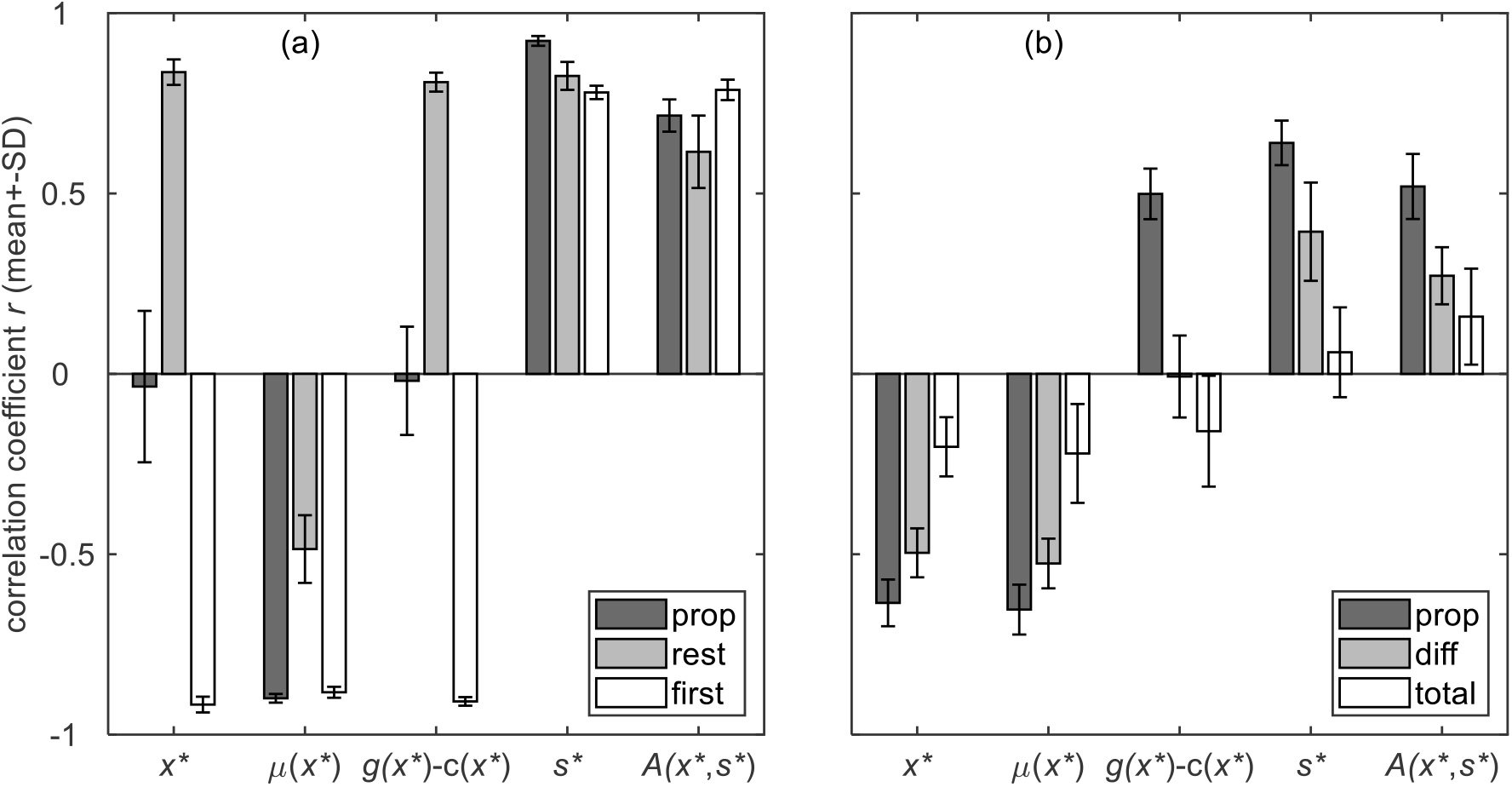
Correlation coefficients between possible proxies (x-axes) and individual fitness (*w*)*. The bars show the *r*^*2*^ (mean ± SD) from linear regressions between each proxy and *w\** for the data shown in Figures 4 and 5, but where variation was added to the proxy and *w\** by adding normally distributed random numbers with standard deviation of 10% of the mean values across the range of environmental variable: (a) predation risk or (b) temperature. All regressions used z-scores of the randomly altered proxy and the randomly altered *w\**. The legend indicates the type of change: proportional increases when foraging and resting (“prop”); constant when resting (“rest); increase in risk first when foraging then resting (“first”); constant difference in cost between foraging and resting (“diff”); constant total cost of foraging (“total”).

To predict the impact of increasing predation risk it is essential to know the specific relationship between foraging rate and predation rate or at least some information about how it is altered by an increase in predator density or a reduced ability to avoid predators. In section 2 I have shown the 3 parameters that should be estimated and how they change with the predation risk. It is important to know whether an increase in the predation risk increases the mortality rate when inactive (ϕ) and how rate increases as activity increases. Studies that measure mortality rates could try to measure the activity rates of individuals and how long they survive. There is a need to agree on a conceptual framework for measuring predation risk (Moll et al., 2017), and I hope that clear theory on the interactions between foraging and mortality will help this process.

Crucial for the mortality rate when resting is the presence of safe refuges and how safe they are. This will depend on environmental complexity, but also on the type of predators. Large predatory fish, for example, will not be able to attack reef fish in amongst the coral, whereas octopuses may be able to (Heithaus et al., 2009). This affects whether ϕ increases. It is also important to know the relationship between an environmental change and both the active metabolic rate and the resting metabolic rate, which could be estimated from similar species, and some information about how these have been affected by the environmental change.

Most studies have assumed that animals have to be active to get any food at all (Abrams, 1984; Boukal, 2014; Houston & McNamara, 1989; Zimmer et al., 2011), but many animals that eat mobile prey can decide whether hunt actively or sit-and-wait (Caraco & Glllespie, 1986; G. Helfman, 1990; G. S. Helfman & Winkelman, 1991; Heller & Milinski, 1979). The models presented here and previous work (Higginson & Ruxton, 2015) show that the availability of food when inactive can strong affect the magnitude and direction of the change in growth and mortality rates if the predation risk increases. If prey animals have a choice of how to catch food then it is crucial to know something of the relative gain rates in order to understand the effect of environmental changes on population viability.

The models here are as simple as possible to review the theory clearly, but many complexities have been omitted. Firstly, in many cases there may be an trait of the animal that alters over time that affects their behaviour (Houston & McNamara, 2014). The most obvious in growing animals is size. The results can equally be used to predict the change in behaviour over the growth period if the size of the animal affects its gain rate, metabolic costs or predation risk. As shown in Figures 4 and 5, if multiple parameters change simultaneously there are many possible outcome patterns. Therefore, researchers could put in the known values or patterns for any particular species into the equation for optimal foraging rate to get predictions for their particular model species before carrying out observations or experiments. I have provided a Matlab function so that anyone can do this.

States other than size may vary that interact with behaviour and life history decisions. Higginson & Ruxton (2010) studied growth under predation risk when an animal can invest in defences. They showed that the relative safety of refuges strongly affects the target size when size affects the risk, and that the effect on foraging rate was slight, which suggests that if animals have flexible maturation size and/or age and can alter some state (e.g. morphological or chemical defence) then they may not respond to a change in predation risk as predicted by simpler models.

Also omitted from the models herein are variability in a the environmental variables. For instance, predation risk may fluctuates over time and animals may choose to forage when the risk is smaller (Lima & Bednekoff, 1999), so it may be important when considering an increase in overall risk how exactly such an increase is distributed. This is all the more important if there is a variable intrinsic state that is under the control of the organism, such as fat reserves that the animal can use to survive periods when it is not profitable and or safe to forage (Higginson et al., 2012).

I have assumed herein that animals can respond appropriately to anthropogenic change. Many studies, including theoretical approaches, have sought to understand how novel predators or foods may cause maladaptive responses by prey (Ehlman et al., 2019; Robertson et al., 2013). One reason that prey animals make counter-intuitive decisions is that they are limited in their behavioural plasticity (Fawcett et al., 2014; Sih et al., 2011), so animals may not be able to perfectly adapt as predicted to an increase in risk if it is too far beyond what they have evolved to cope with. The development of mechanistic models of the physiological and psychological control of behaviour will be useful for predicting prey responses (Giske et al., 2013; Gross et al., 2008; Higginson et al., 2016), as these will predict the extent of behavioural plasticity.

It will also be important to predict the behavioural changes of more than just one species in the interaction. For instance, predators may be less effective under noise (Barton et al., 2018), which may dampen the increase in prey vulnerability. Richer game theoretic models are necessary to predict these outcomes (McNamara, 2013). A research programme based on synergistic interplay between theoretical and empirical work will greatly help researchers understand and thereby mitigate the impacts of anthropogenic changes on ecosystems.

## Supporting information

Matlab function (after download change extension to .m)

## ACKNOWLEDGEMENTS

The author was supported a NERC Independent Research Fellowship (NE/L011921/1).

---

**Table 1:** Parameters in the models and their default values

| Parameter / Variable | Symbol | Baseline | Values |
| --- | --- | --- | --- |
| Ratio of mortality to gain rate | $v$ | Derived | – |
| Rate of predation | $\mu$ | Derived | – |
| Rate of energy gain | $\gamma$ | Derived | – |
| Rate of energy use | $c$ | Derived | – |
| Rate of foraging | $x$ | Optimised | – |
| Activity-associated predation rate | $p$ | 0.5 | |
| Baseline mortality rate | $\phi$ | 0.1 | |
| Activity-associated gain rate | $r$ | 1.0 | |
| Relative gain rate when inactive | $\alpha$ | 0.1 | |
| Activity-associated metabolic cost | $m$ | 0.2 | |
| Relative metabolic cost when inactive | $\beta$ | 0.5 | |

