## Appendices for "Identifying reliable fitness proxies for growing animals responding to anthropogenic changes"

**APPENDIX A: ORIGIN OF ACCELERATING EFFECT OF FORAGING ON PREDATION RISK**

Models of foraging often assume that mortality rate is an accelerating function of foraging intensity, such that low levels of foraging are not very costly but maximising foraging rate results in very high mortality (Houston et al., 1993; Werner & Gilliam, 1984). This is an assumption that results in solutions to the equations because it means that maximum foraging is prohibitively costly, and this suggests it is the case in nature. Here, I show one situation in which it is likely to be case: vigilance for an attack by a predator that can be avoided if detected.

Consider a situation where it takes $\tau$ seconds for a predator to successful catch a prey from the time it can be detected (e.g. leaving cover). The prey animal has to decide what proportion of time to be vigilant (e.g. scan with raised head) versus eating (head lowered). Assume that scans are randomly dispersed over time and occur with probability $q$ . An attack occurs with frequency $p$ and will be successful if a scan does not occur for the $\tau$ seconds it takes for the predator to reach the prey from cover. The probability that a random scan doesn't occur during this time is therefore binomially distributed with number of trials $\tau$, success probability $q$ and no successes ($k$=0).

$$\Pr(successful\ attack) = (1-q)^{\tau} \tag{A1}$$

Since $0 \leq q \leq 1$ the part in parentheses is a proportion $0 \leq q \leq 1$ so this function is decelerating. Since the amount of time spent foraging is $x = 1 - q$, this is accelerating with respect to foraging rate: $x^{\tau}$ (Figure A1a). Consider the case where the gain from foraging is constant, so just $x$. Then the mortality to growth rate assuming rate of attack $p$ and some other source of mortality $\phi$ is

$$v = \frac{px^{\tau} + \phi}{x} \tag{A2}$$

shown in Figure A1b. And the optimal proportion of time that the animals is foraging rather than vigilant is given by

$$x^* = \left(\frac{\phi}{p[\tau - 1]}\right)^{\frac{1}{\tau}} \tag{A3}$$

Note that if $\phi < p$ and $\tau \geq 2$ the part in parentheses is a proportion $0 \leq q \leq 1$ so $x^*$ increases as $\tau$ increases. That is, the longer time the animal has to react to an attack, the more time they can spend foraging. In the main text, $\sigma$ has a similar effect to $\tau$: it also alters the shape of the relationship and so the mortality at intermediate foraging rates ($x$).

**Figure A1**: (a) Mortality rate as a function of foraging effect *x* for four values of the duration of time an imminent attack can be detected ($\tau$). (b) The mortality to growth ratio ($v$) for the same values of *x* and $\tau$. Other parameter values: $p$=0.6, $\phi$=0.1. The optimal *x* is that which minimises $v$.

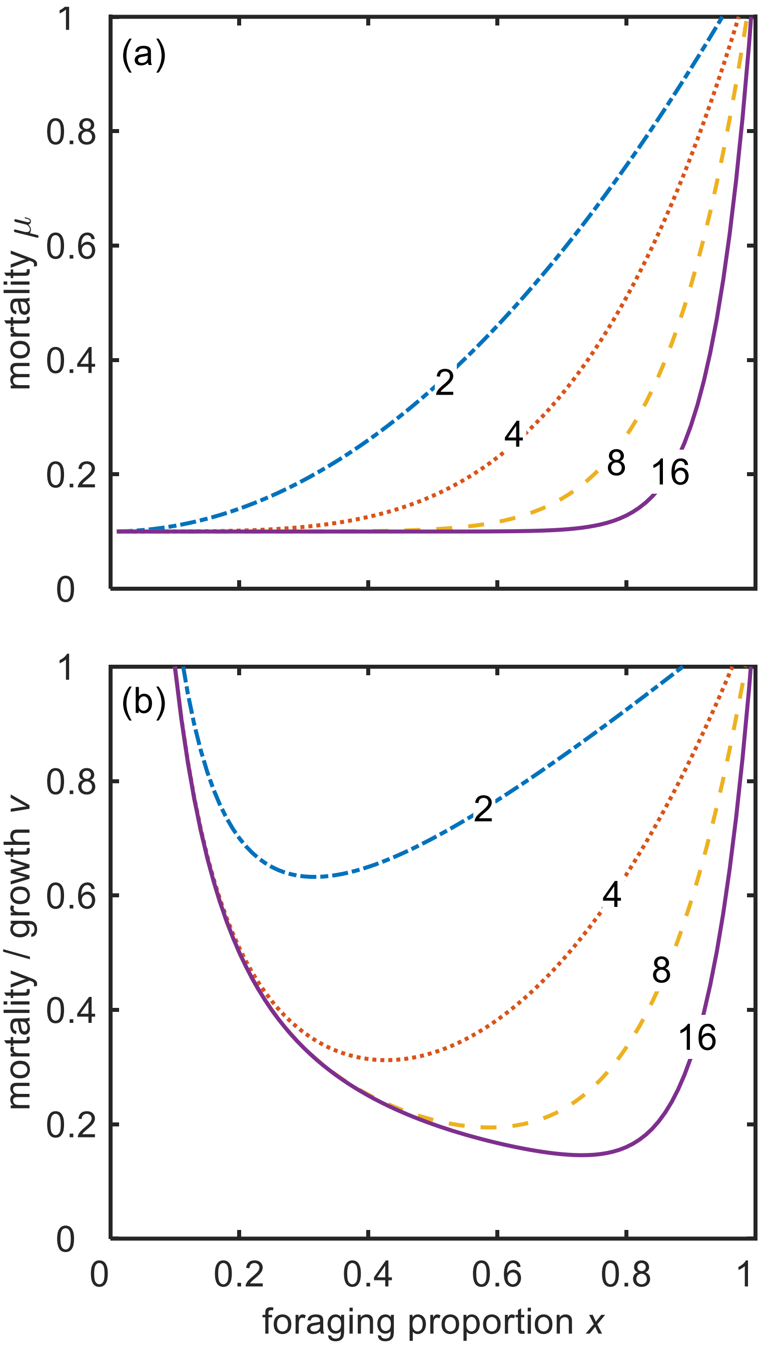

### APPENDIX B: GENERAL METHOD FOR PREDICTING MATURATION

The mortality to growth ratio can be used to predict a choice in any given state. This approach can be extended to take into account the value of being in any given state when the animal must trade off the benefit of further growth against the risk of dying and so losing everything (Houston & McNamara, 1989). One application is to predict the optimal timing of maturation for an animal that gets a greater reproduction if larger but when growing carries a risk of mortality (English et al., 2016),(Appendix B). Effectively, there is a trade-off between the increasing payoff after maturity and the probability of reaching maturity (Figure B1a). The animal should forage more to grow rather than mature if the following inequality holds

$$\frac{ds(x)}{dt}\frac{dw}{ds} - \mu(x)w(s) > 0 \qquad \text{(B1)}$$

where $w$ is the current expected reproductive success for size $s$, μ is the probability of death, $\frac{ds}{dt}$ is the rate of increase in size against time, and $\frac{dw}{ds}$ is the rate of increase in $w$ given an increase in size. The first term (positive) decreases as size increases as the effect of body size on reproduction has diminishing returns (Figure B1b). The second term (negative) increase with body size as the reproductive potential gets larger so there is more to lose by being killed. At the point at which they are equal, the animal should stop growing and mature. The optimal foraging rate found from solving for $x$ in the mortality to growth ratio $v$ gives us the optimal growth rate $\frac{ds}{dt}$ and the optimal mortality rate μ. If there is no effect of size on gain or mortality then the optimal foraging rate is the same for all sizes. Thus, for any assumed function of $w(s)$ we can use this formation to find how size at maturation interacts with optimal foraging rate and how they depend on predation and metabolic costs.

**Figure B1**: Illustration of the use of inequality B1 to predict optimal size at maturation. (a) The reproductive payoff (solid line) from body size *s* (x-axis) had diminishing returns, here $w = \sqrt{s - \frac{1}{2}}$ . Given a constant foraging rate the proportion of individuals still alive (dashed line) declines with a constant proportion from unity. (b) The positive term of inequality B1 is initially large because small adults have very little reproductive value, and levels off. The negative term increases as reproductive potential increases (dotted line). When the sum of these (dashed line) is zero, the animal should mature. Plots are based on the equations in the main text with the baseline parameters values (Table 1).

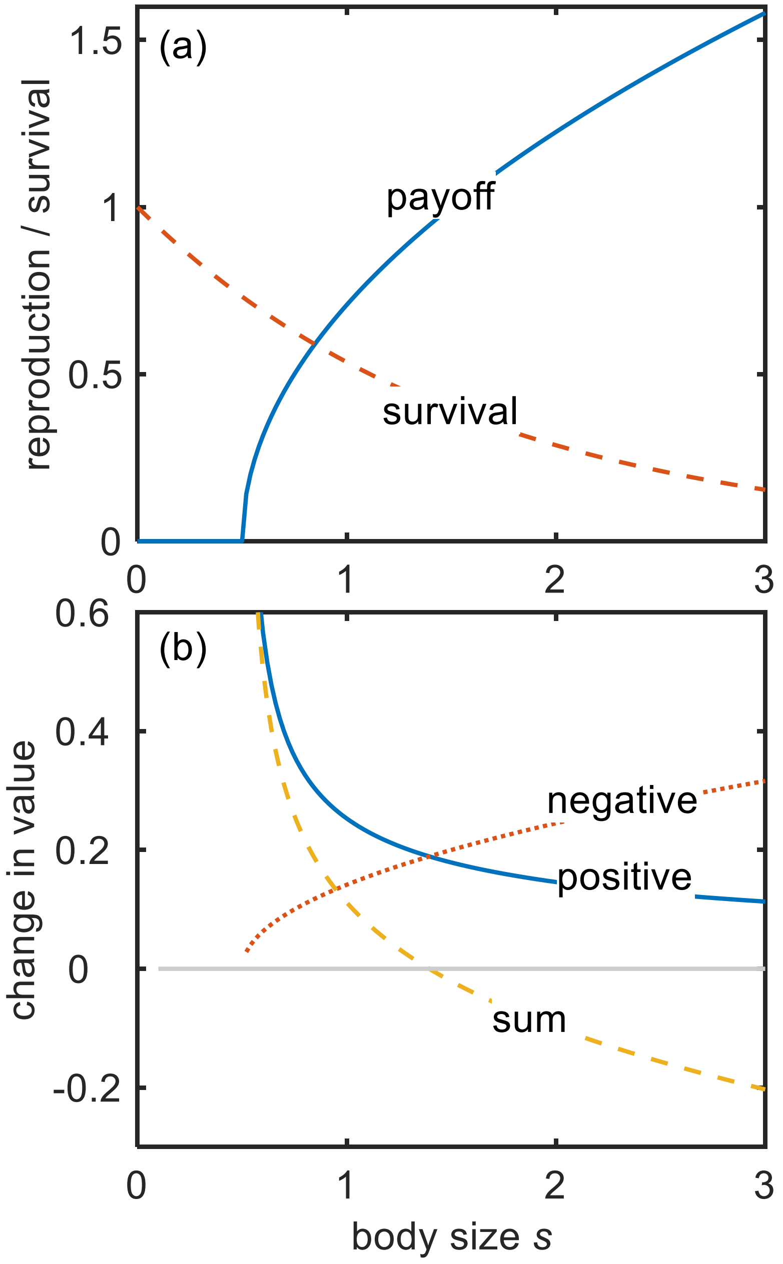

### APPENDIX C: SUPPLEMENTARY FIGURES

**Figure C1**: Effect of varying each parameter (columns) in turn from baseline values (*r*=1.0, α=0.1, *m*=0.2, β=0.5, *p*=0.5, ϕ=0.1). Top row: effect of foraging *x* on the mortality to growth ratio *v* for three values of the parameter (shown on lines). Middle row: optimal foraging rate *x**, Bottom row: optimal gain *g** (dashed lines) and mortality rate *μ** (solid lines).

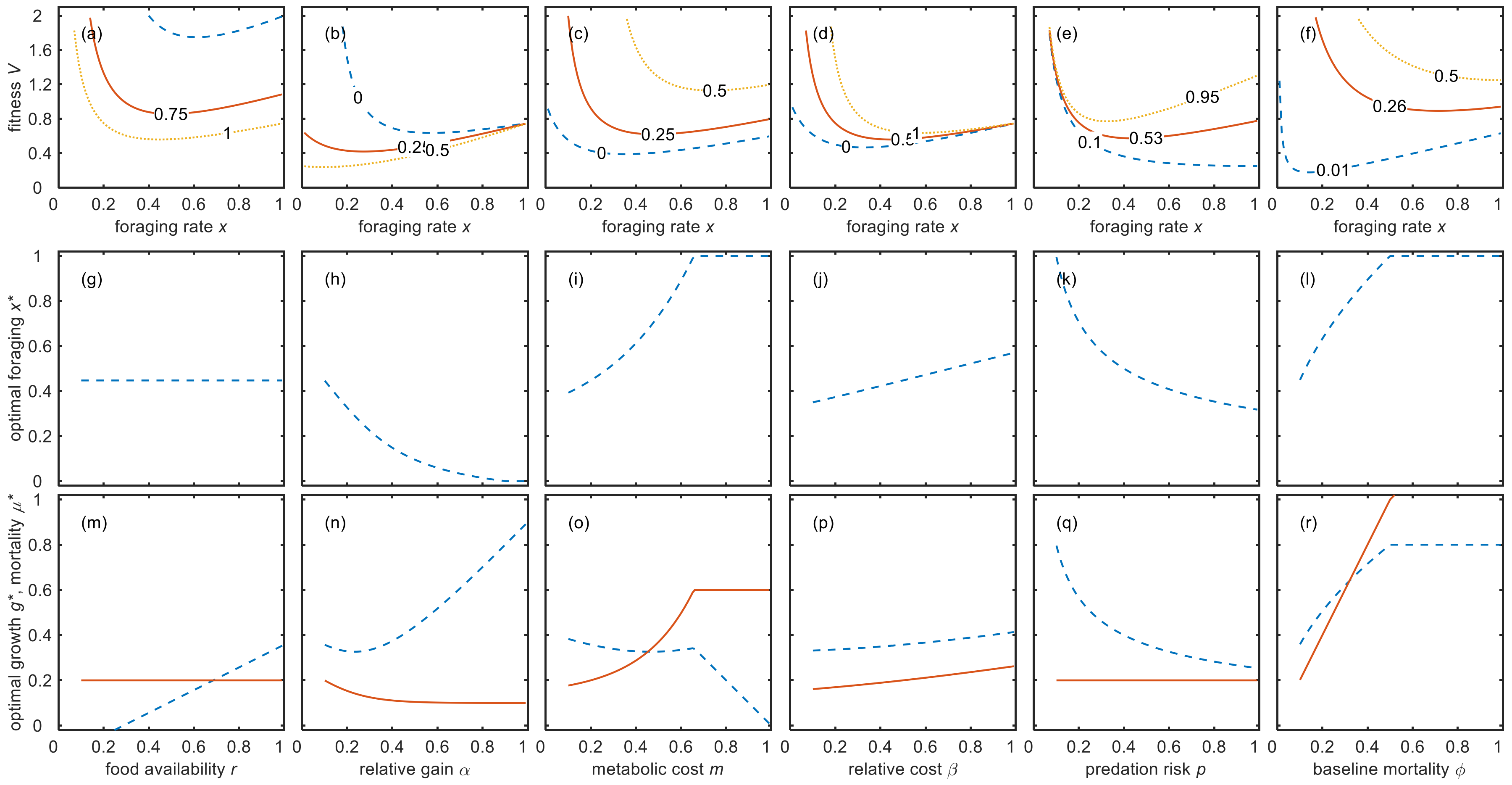

**Figure C2**: Effect of each parameter (columns) on the optimal foraging rate (green negative; cyan negligible; blue positive) for the values of the parameter (x-axes) and the other parameters (rows). Black areas are where $x^*\geq 1$ and white areas where $x^*\leq 0$ so does not change when the parameter changes by a small amount. These areas are also unfeasible for most situations because, for example, costs are almost the same as the maximum gain (right of 2$^{nd}$ column) or the maximum predation rate is smaller than the baseline mortality rate (right of 4$^{th}$ column).

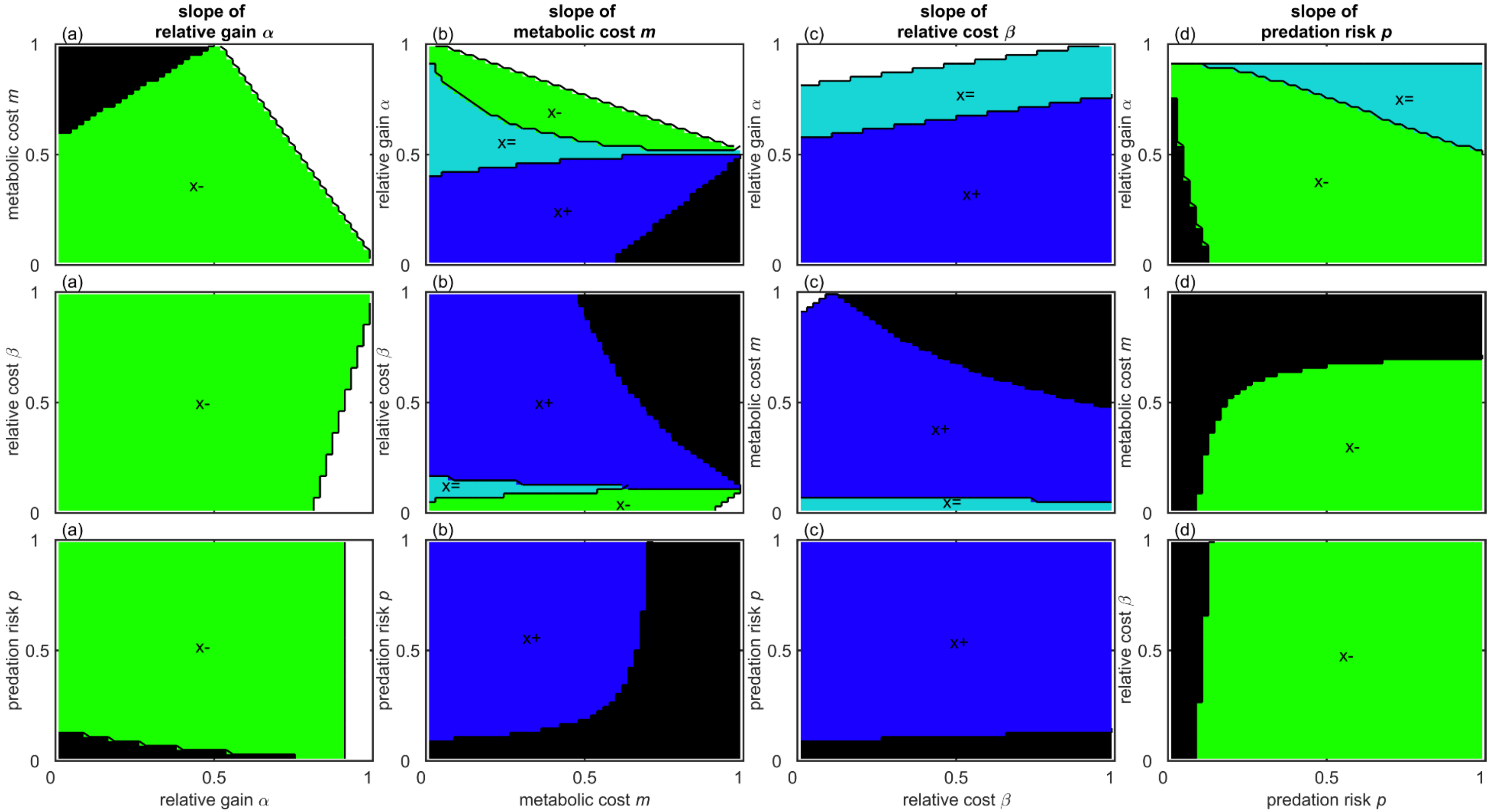

**Figure C3:** Effect of each parameter (columns) on the growth and mortality rates when foraging is optimised for the values of the parameter (x-axes) and the other parameters (rows, shown on y-axes). Black areas are where $x^*\geq 1$ and white areas where $x^*\leq 0$ so does not change. Effects (positive +, negative −, negligible =) are shown for growth rate (g) and mortality rate (μ) in labels in coloured areas.

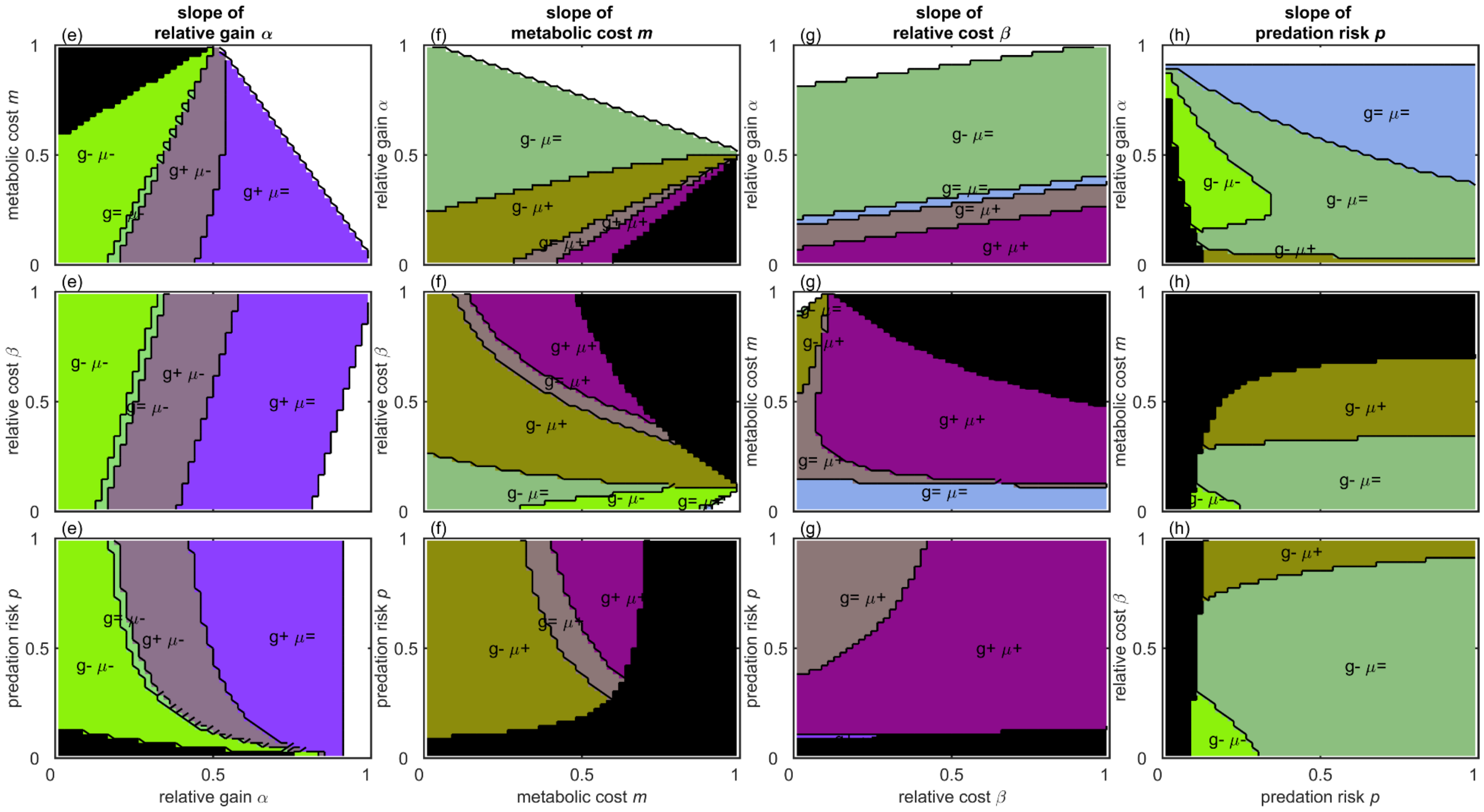

**Figure C4:** Effect of each parameter (columns) on size at and survival to maturation when foraging is optimised for the values of the parameter (x-axes) and the other parameters (rows, shown on y-axes). Black areas are where $x^{*} \geq 1$ and white areas where $x^{*} \leq 0$ so does not change. Effects (positive +, negative −, negligible =) are shown for size at maturation ($s$) and survival to maturation ($A$) in labels in coloured areas.

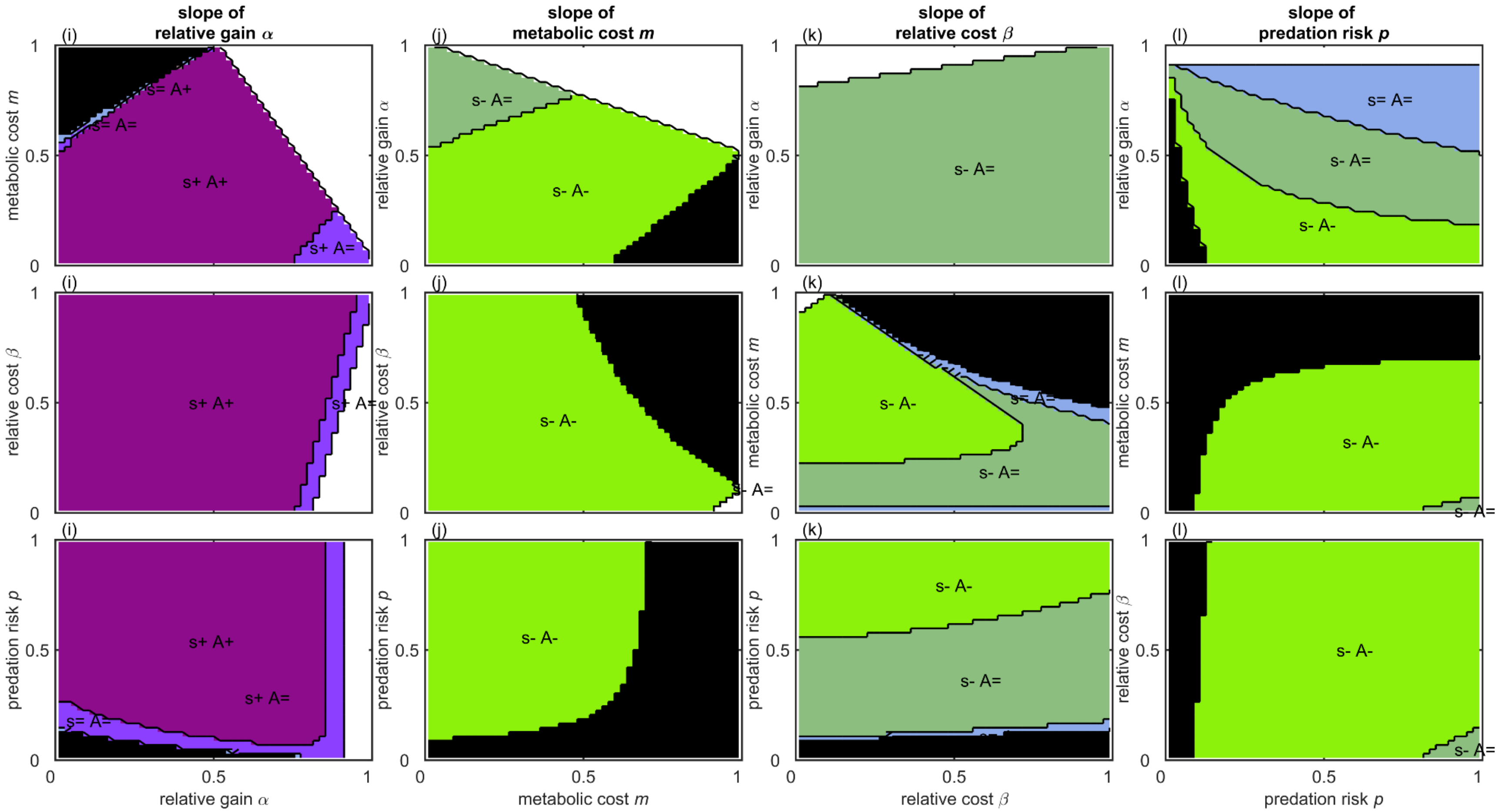

**Figure C5**: Environmental changes or experiments that increase predation risk $q$ (e.g. predator density) or energy use could affect the parameters in at least 3 ways. The lines show the mortality rate (a, b, c) and energy use (d, e, f) from foraging (dotted lines: $p$ or $m\beta$) and from resting (dashed lines) and the total (solid lines; which the mortality or energy use when maximally foraging). (a) Increase risk both when foraging and not foraging: $p = kq$ and $\phi = (1-k)q$, where $k$ is a positive constant $k = 2/3$, $0 \le q \le 1$. (b) Increase risk only when foraging: $p = (1-k)q$ and $\phi = k$, $k = 1/5$; (c) Increasing risk first increasing risk when foraging and then for resting because e.g. predators avoid one another so there are fewer safe places $p = q - (1-k)q^2$ and $\phi = (1-k)q^2$, $k = 2/3$. (d) Increasing energy use equally when foraging and resting as ambient temperature $q$ increases $m = \frac{q}{2}$ and $\beta = k$, $k = 1/2$; (e) Constant cost of foraging and increasing baseline metabolic rate $m = \frac{k+q}{4}$ and $\beta = \frac{q}{k+q}$ $k = 6/10$; (f) Constant total cost of foraging, decreasing baseline metabolic rate matched by increasing cost of foraging $m = k$ and $\beta = 1 - q$, $k = 3/10$.

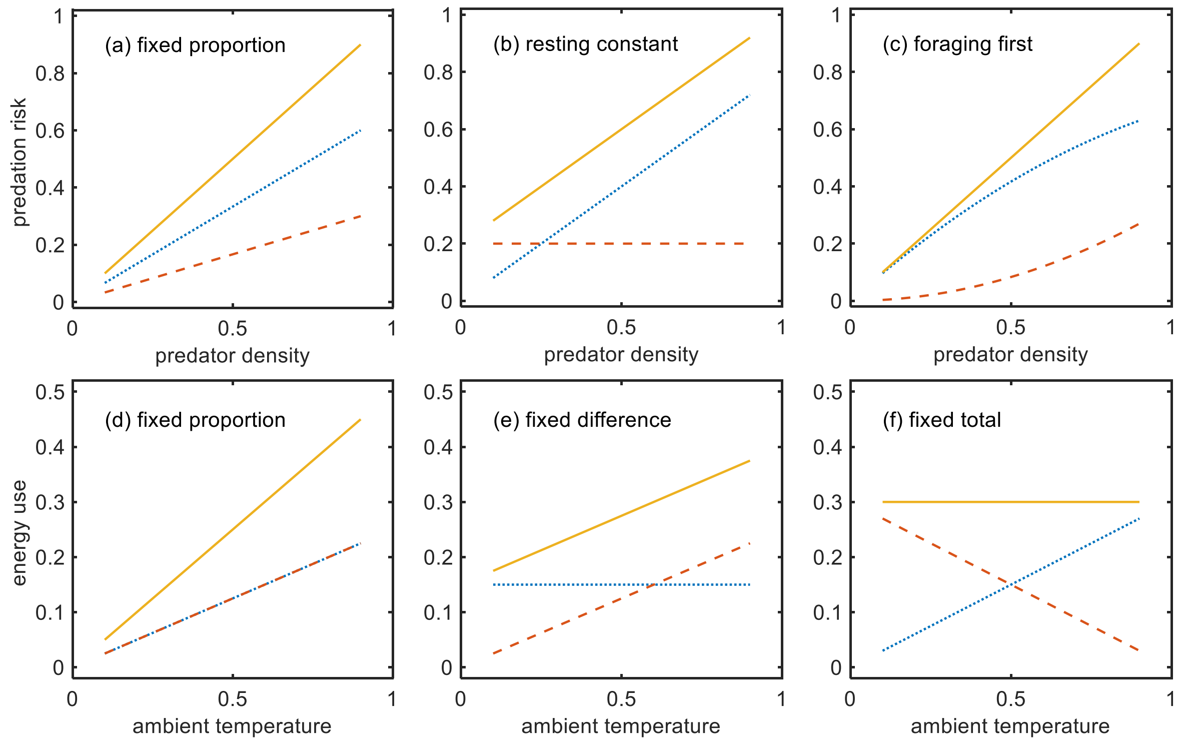

**APPENDIX D: CONDITIONS FOR CHANGES IN RESPONSES**

Here, I show how the growth, mortality, size at and survival to maturation change when the parameters change. The effects are presented in Figures 3 and C2-4.

**Growth and mortality**

The optimal foraging rate is

$$x^* = \frac{m\beta - \alpha}{1 - \alpha - m + m\beta} \pm \frac{\sqrt{\alpha^2 - 2\alpha m\beta + m^2\beta^2 + \phi[(1-m)^2 + (1-m)(2m\beta - 2\alpha)]}}{\sqrt{1-\phi}(1 - \alpha - m + m\beta)} \qquad (D1)$$

This expression is highly complex. For some intuition consider the case where that there is no gain when not foraging (α=0) and the activity costs are small enough that they can be ignored (*β*=1). Then the optimal foraging rate over all of the juvenile period is

$$x^* = m \pm \sqrt{m^2 + \frac{\phi}{p}} \qquad (4.1)$$

so the foraging rate increases if *m* or $\phi$ increase, and decrease if *p* increases. The subsequent mortality rate is

$$\mu(x^*) = \phi + p\left(m + \sqrt{m^2 + \frac{\phi}{p}}\right)^2 \qquad (4.2)$$

and the optimal growth rate is

$$g(x^*) - c(x^*) = \sqrt{m^2 + \frac{\phi}{p}} \qquad (4.3)$$

The changes in *x** are always as expected, except for when *m* increases if β < α, when an increase in *m* causes *x** to decrease due to the different relative costs. *x** is greater than unity if

$$\alpha > \frac{\phi(1 - m + m\beta) - p(1 - m - m\beta)}{\phi + p}$$

so the foraging rate will not change in this parameter space for small changes in parameter values.

The change in growth rate with an increase in the availability of free food is

$$\frac{d[g(x^*) - c(x^*)]}{d\alpha} \qquad (D2)$$

and so solving this equal to zero gives the condition for an increase in growth rate when free food becomes available.

$$\alpha > m\beta + \frac{\phi(1-m)}{\phi + p} \qquad (D3)$$

Foraging rate always declines if α increases, so if α is sufficiently high (not foraging has high gains) then growth increases. This depends on the metabolic costs and predation rate when not foraging because these affect the extent of the change in foraging rate. Secondly, the change in mortality rate with an increase in the availability of free food is

$$\frac{d\mu(x^*)}{d\alpha} \tag{D4}$$

and so solving this equal to zero gives the critical value of α above which mortality rate increases when free food availability increases. There is no solution for this, so free food always decreases mortality rate.

Next I consider an increase in *m*, which could be thought about as either an increase in overall metabolic costs or as a reduction in overall food availability. Growth increases if *m* increases under the following condition

$$m > \frac{\alpha\beta(\phi + p) + \phi(1 - \alpha - \beta)}{\beta^2(\phi + p) + \phi(1 - 2\beta)} \tag{D5}$$

There is an increase in mortality if $\beta < \alpha$, which could occur if activity is very expensive and/or gain is high when inactive, such as for a specialised poikilothermic sit-and-wait predator.

Next I consider the effect of an increase in β, which might occur if the temperature increases and the animal is an ectotherm or the animal is an endotherm in cold conditions so they need to spend energy staying warm when not active. Mortality always increases in this case, but growth increases only if

$$\beta > \frac{\alpha(\phi + p) - \phi(1 - m)}{m(\phi + p)} \tag{D6}$$

due to the increase in foraging activity because the benefit of resting is smaller.
An increase in *p* decreases mortality if

$$\alpha > m\beta \tag{D7}$$

and increases growth if

$$m(1 - \beta) < 1 - \alpha \tag{D8}$$

This occurs because if α is sufficiently large then the foraging rate drops sufficiently to reduce mortality, which is always lower when not foraging. There is never an increase in growth because if α is sufficiently high for 3.13 to be satisfied then $x^*<0$.

An increase in $\phi$ always increases mortality and *increases* growth, as might be expected from section 1 and 2.

**Size at maturation**

Assume that the fitness after maturation *w* is simply the square root of body size *s* minus some minimum size for maturation (ψ),

$$w = \sqrt{s - \psi}$$

so this has diminishing returns, i.e. the positive slope gets smaller as *s* increases,

$$\frac{dw}{ds} = \frac{1}{2\sqrt{s-\psi}} \tag{D9}$$

The inequality from Appendix B is therefore

$$\frac{r\{x+\alpha(1-x)\}\ -m\{x+\beta(1-x)\}}{2\sqrt{s-\psi}} - (px^2+\phi)\sqrt{s-\psi} > 0 \tag{D10}$$

If this is negative the animal should mature. Rearranging this equation gives the critical size for maturation

$$s^* = \psi + \frac{\alpha r - \beta m + x\{r(1-\alpha) - m(1-\beta)\}}{2(px^2+\phi)} \tag{D11}$$

Note that if $x = 1$ then optimal size is

$$s^* = \frac{r-m}{2(px^2+\phi)} \tag{D12}$$

If $x$ does not change the maturation size increase if $r$ increases and decreases if $m$, $p$ or $\phi$ increases. For the optimal foraging rate ($x$*) equation D12 is complex, but in the case where there is no gain when not foraging (α=0) and that the activity costs are small enough that they can be ignored ($\beta$=1).

$$s^* = \psi + \frac{\sqrt{m^2 + \frac{\phi}{p}} - m}{4\phi} \tag{D13}$$

An increase in $p$, ϕ, and $m$ all cause a decrease in maturation size.
By differentiation we can show that $s$* increases if α increases if

$$\frac{p}{\phi} < \frac{2(1-m)}{1+\alpha-m(1+\beta)} - 1 \tag{D14}$$

And increases if $m$ increases if

$$\frac{p}{\phi} < \frac{2(\beta-\alpha)(1-\beta)^2}{\beta^2[\beta-2\alpha+\alpha\beta+m\beta(1-\beta)]} - \frac{(1-\beta)^2}{\beta^2} \tag{D15}$$

And increases as β increases if

$$\frac{p}{\phi} < \frac{2(1-m)}{1+\alpha-m(1+\beta)} - 1 \tag{D16}$$

Which is the same condition as for α. The values of parameters where the opposite is predicted are unfeasible, and where $x$*>1.

**Survival to maturation**

The time to maturation, assuming that growth is constant, is proportional to the difference between the size at maturation and the size at hatching/birth divided by the growth rate. The survival to maturation is the rate of survival to the power of the time. It doesn't matter what value we give to size at hatching/birth so assume it is a zero, then the proportion of individuals that mature is

$$A = e^{-\mu(x^*)\frac{s^*}{g(x^*)-c(x^*)}} \tag{D17}$$

If there is a minimum size then

$$-\log(A) = (px^2 + \phi\,)\frac{\psi + \dfrac{f - m - (1-x)\{f(1-\alpha) - m(1-\beta)\}}{2(px^2+\phi)}}{r\{x + \alpha(1-x)\} - m\{x + \beta(1-x)\}} \qquad (D18)$$

Which simplifies to

$$-\log(A) = \frac{\psi(px^2 + \phi\,)}{r(x + \alpha\{1-x\}) - m(x + \beta\{1-x\})} + \frac{1}{2} \qquad (D19)$$

Note that if there is no minimum size (ψ=0) then the survival to maturation is a constant

$$A = e^{-\frac{1}{2}} = 0.607 \qquad (D20)$$

The effect of each parameter on survival to maturation hence varies with the parameters, from very small to large.

### APPENDIX E: ADAPTIVE CHANGES IN METABOLISM

Growing animals have to meet basic metabolic needs but sometimes show elevated metabolic rate (as measured by oxygen consumption), which is interpreted as stress. On some occasions animals may adjust their metabolism to the environment (Careau et al., 2008; Le Galliard et al., 2013; Preisser & Orrock, 2012). Models of foraging that assume that foraging rate and metabolism can be used to predict optimal responses to changes in predator density or food use (Houston 2010).

Assume that the predation risk is the ratio of the encounter rate with predators to the metabolism, so that vulnerability declines with investment in metabolism but with diminishing returns.

$$\mu = \frac{px^2}{m} + \phi \qquad (E1)$$

The gain rate is as the main text, but with no gain when not foraging (α=0).

$$g = rx \qquad (E2)$$

The metabolic cost is the baseline metabolic cost (β) plus the extra cost of foraging

$$c = \beta + mx$$

As before, we assume the animal minimises the ratio of the mortality to growth rates

$$v = \frac{\frac{px^2}{m} + \phi}{rx - mx - \beta} \qquad (E3)$$

The optimal foraging rate is

$$x^* = \frac{\beta p \pm \sqrt{p(p\beta^2 + \phi r^2 m - 2\phi r m^2 + \phi m^3)}}{p(r - m)} \qquad (E4)$$

The negative root always gives a negative result (i.e. impossible) for reasonable values of the parameters. Since the terms under the square root must sum to a positive value if the result is to be real (not imaginary) and *m* appears in the denominator with a negative sign, the positive root always increases as *m* increases (Figure E1, black line). For example, note that when ϕ=0 (i.e. other mortality risks can be ignored) the positive root is

$$x^* = \frac{2\beta}{r - m} \qquad (E5)$$

The optimal metabolic rate also increases as foraging rate increases

$$m^* = \frac{px^2 \pm \sqrt{px(px^3 + \phi rx - \beta\phi)}}{\phi} \qquad (E6)$$

Again, all the *x* terms are positive so *m** increases if *x* increases (Figure E1, white line).
By substituting one into the other we can find the optimal combination (intersection of white and black lines in Figure E1a). Surprisingly, the response in foraging rate results in the metabolic rate depending only on the food availability

$$m^*(x^*) = \frac{r}{3} \qquad (E7)$$

$$x^*(m^*) = \frac{9\beta p \pm \sqrt{81p^2\beta^2 + 12\phi r^3 p}}{6pr} \qquad (E8)$$

Whilst when ϕ=0 is

$$x^*(m^*) = \frac{3\beta}{r} \qquad (E9)$$

In Figure E2 I also consider what happens when foraging cannot be altered. This may occur if in an experiment the animals cannot alter its activity pattern (e.g. in a swim tunnel respirometer) or if the foraging rate is already maximal ($x$*=1). If food availability increases then foraging decreases and energy use increases (Figure E2a). As the baseline costs increase (Figure E2b) foraging rate would increase and energy use would be constant, up to the point where foraging could not increase any further and costs have to be reduced, even though this increases vulnerability to predation. If predation risk increases the foraging effort declines, but if it cannot then energy use increases to help avoid predator attacks. The opposite is predicted if the other source or mortality increases.

**Figure E1**: Mortality to growth ratio as a function of the foraging rate (*x*) and metabolic rate (*m*) for the baseline parameter values (*r*=1.0, β=0.2, *p*=0.5, ϕ=0.1). (a) Landscape in (*x*, *m*) space and colours showing *v* (see sidebar). White indicates where *v* < 0 because of insufficient foraging and too high cots. (b) Cross sections through the landscape at 4 values of *x* showing how in the region around the optimal strategy (*x**,*m**) the landscape is quite flat, indicating that a range of strategies have roughly equal fitness.

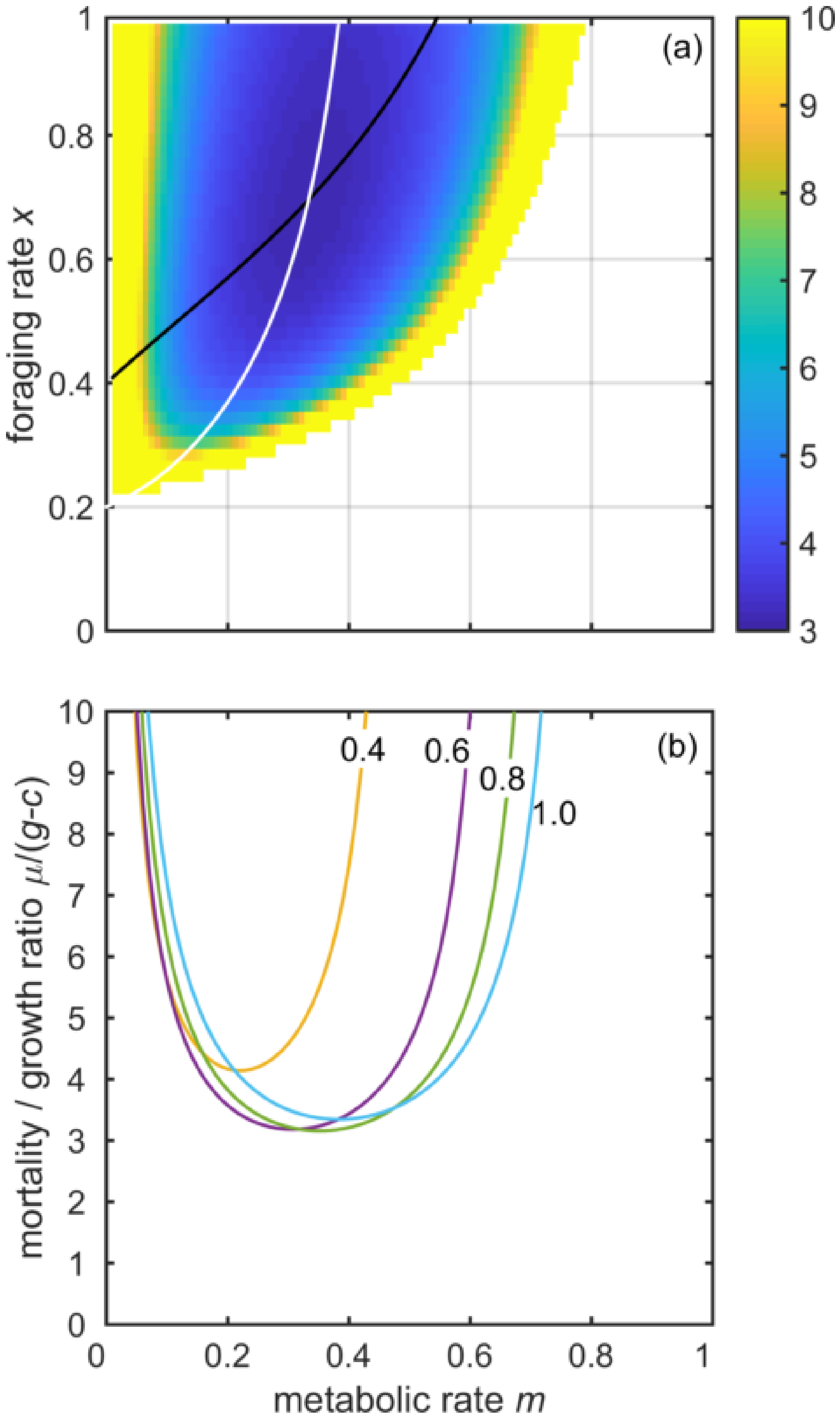

**Figure E2**: Optimal strategies when foraging rate and metabolic rate can both be adjusted. Each panel shows the effect of different parameters: (a) food availability, (b) baseline metabolic cost β, (c) predation risk *p*, (d) baseline mortality *ϕ*. There are four lines on each panel: optimal foraging rate *x** (grey lines), optimal metabolic rate when optimal foraging rate is possible [*m**(*x**) solid black lines], optimal metabolic rate when foraging rate is fixed at the baseline value [*m**(*x*) dashed lines], optimal metabolic rate when foraging rate is greater than unity so suboptimal at unity [*m**(*x*=1) dotted lines].

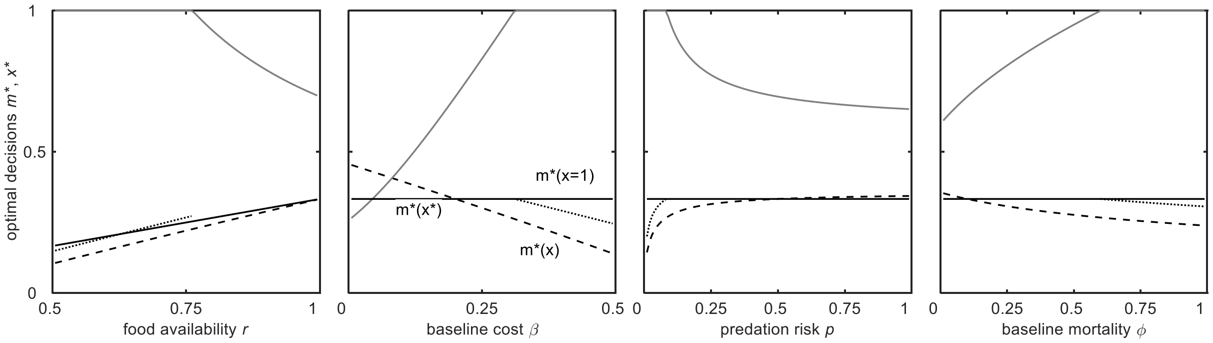
