## Supplementary material for "Identifying reliable fitness proxies for growing animals responding to anthropogenic changes": Matlab function (after download change extension to .m)

function [optxx,optmort,optgrowth,optsize,optsurvival,optfitness]=growthrisk_yourpredictions(varargin)
% GROWTHRISK_YOURPREDICTIONS generates predictions based on Higginson (2020) 
%      "Identifying reliable fitness proxies for growing animals responding to anthropogenic changes"
% 
%      [OPTXX,OPTMORT,OPTGROWTH,OPTSIZE,OPTSURVIVAL,OPTFITNESS]=growthrisk_yourpredictions(basepred,halfpred,maxpred,amr,bmrprop,freefood)
%      input are name-value pairs with vectors of any length (including 1x1) but any that
%      are not scalars (1x1) must be equal size. 
%              basepred: baseline mortality rate independent of foraging (default 0.1)
%              halfpred: mortality rate at half foraging rate (default 0.225, so sigma=2)
%              maxpred: mortality rate at maximum foraging rate (default 0.6)
%              amr: relative energy use compared to maximum gain rate (default 0.2)
%              bmrprop: proportional use of energy when resting compared to when foraging (default 0.4)
%              freefood: proportion gain when resting compared for when foraging (default 0.2)
% 			
% 		       example 1: effect of an increase in energy use
% 			   [...]=growthrisk_yourpredictions('amr',[0.1 0.2]);  
%              example 2: effect of a doubling in predation risk for all levels of foraging rate
%              [...]=growthrisk_yourpredictions('maxpred',[0.5 1.0],'halfpred',[0.22 0.44],'basepred',[0.1 0.2])
%              example 3: effect of gradually increasing BMR as proportion
%              of AMR from negligible to equal
%              [...]=growthrisk_yourpredictions('bmrprop',0:0.05:1)
%
%      outputs for each value of the input vectors:
%              OPTXX optimal foraging rate
%              OPTMORT mortality rate
%              OPTGROWTH growth rate
%              OPTSIZE size at maturation
%              OPTSURVIVAL survival to maturation
%              OPTFITNESS fitness = OPTSURVIVAL * (value of  OPTSIZE)
% 
% (c) Andrew D. Higginson 06/08/2020

% define dafaults 
options = struct('basepred',0.1,'halfpred',0.225,'maxpred',0.6,'amr',0.2,'bmrprop',0.4,'freefood',0.2);
% read the acceptable names
optionNames = fieldnames(options);
% count arguments
nArgs = length(varargin);
if round(nArgs/2)~=nArgs/2
   error('inputs need to be name/value pairs')
end
for pair = reshape(varargin,2,[]) %# pair is {propName;propValue}
   inpName = lower(pair{1}); %# make case insensitive
   if any(strcmp(inpName,optionNames))
      options.(inpName) = pair{2};      
   else
      error('%s is not a recognized parameter name',inpName)
   end
end
% extract variables from structure 
basepred=options.basepred;
halfpred=options.halfpred;
maxpred=options.maxpred;
amr=options.amr;
bmrprop=options.bmrprop;
freefood=options.freefood;
% number of foraging values
nx=10001;
% number of settings
nset=max([numel(basepred),numel(halfpred),numel(maxpred),numel(amr),numel(bmrprop),numel(freefood)]);
% numerical
xx=repmat(linspace(0,1,nx),[nset 1]);
% make all inputs NSET x length(xx)
basepred=makesame(basepred,nx,nset);
halfpred=makesame(halfpred,nx,nset);
maxpred=makesame(maxpred,nx,nset);
amr=makesame(amr,nx,nset);
bmrprop=makesame(bmrprop,nx,nset);
freefood=makesame(freefood,nx,nset);
% work out what sig is from the user input
pred=maxpred-basepred;
% by rearragement of mu=phi+pred.*xx^sig for xx=1./2
sig=log((halfpred-basepred)./pred)./log(1/2);
% mortality rate
mort=pred.*xx.^sig+basepred;
% energy gain rate
gane=(xx+freefood.*(1-xx));
% energy use rate
cost=amr.*(xx+bmrprop.*(1-xx));
vv=mort./(gane-cost);
% don't allow spurious solution
vv(gane<cost)=max(vv(:));
% find index of lowest v
[~,xind]=min(vv,[],2);
% get foraging rate 
optxx=xx(1,xind);
% get mortality and growth rates
optmort=mort(1:nset,xind);
optmort=diag(optmort)';
optgrowth=gane(1:nset,xind)-cost(1,xind);
optgrowth=diag(optgrowth)';
% when to mature
minSize=0.5;
bsize=repmat(linspace(minSize+10^-6,5,nx),[nset 1]);
% fitness at maturation
ww=(bsize-minSize).^(1/2);
dwds=1./(2.*(bsize - minSize).^(1/2));
% mcnouston equation
w2=repmat(optgrowth',[1 nx]).*dwds-repmat(optmort',[1 nx]).*ww;
% size of maturation
[~,sizeind]=min(abs(w2),[],2);
optsize=bsize(1,sizeind);
% survival to maturation
optsurvival=exp((-1.*optmort).*(optsize./optgrowth));
% fitness of population
optfitness=ww(1,sizeind).*optsurvival;
